# DNA-PKcs Suppresses Illegitimate Chromosome Rearrangements

**DOI:** 10.1101/2023.12.30.573736

**Authors:** Jinglong Wang, Cheyenne A Sadeghi, Richard L Frock

## Abstract

Two DNA repair pathways, non-homologous end joining (NHEJ) and alternative end joining (A-EJ), are involved in V(D)J recombination and chromosome translocation. Previous studies reported distinct repair mechanisms for chromosome translocation, with NHEJ involved in human and A-EJ in mice predominantly. NHEJ depends on DNA-PKcs, a critical partner in synapsis formation and downstream component activation. While DNA-PKcs inhibition promotes chromosome translocations harboring microhomologies in mice, its synonymous effect in humans is not known. We find partial DNA-PKcs inhibition in human cells leads to increased translocations and the continued involvement of a dampened NHEJ. In contrast, complete DNA-PKcs inhibition substantially increased microhomology-mediated end joining (MMEJ), thus bridging the two different translocation mechanisms between human and mice. Similar to a previous study on Ku70 deletion, DNA-PKcs deletion in G1/G0-phase mouse progenitor B cell lines, significantly impairs V(D)J recombination and generated higher rates of translocations as a consequence of dysregulated coding and signal end joining. Genetic DNA-PKcs inhibition suppresses NHEJ involvement entirely, with repair phenotypically resembling Ku70-deficient A-EJ. In contrast, we find DNA-PKcs necessary in generating the near-excusive MMEJ associated with Lig4 deficiency. Our study underscores DNA-PKcs in suppressing illegitimate chromosome rearrangement while also contributing to MMEJ in both species.

## Introduction

The catalytic subunit of DNA-dependent protein kinase, DNA-PKcs, is a large protein (∼460 Kd) present in vertebrates, and orthologs of this protein exist in a wide range of organisms including invertebrates, fungi, plants, and protists.^1^ DNA-PKcs forms a DNA-PK holoenzyme complex by binding with Ku70/80 and serves primarily in non-homologous end joining (NHEJ)-mediated repair of DNA double-stranded breaks (DSBs).^2,3,4^ DNA-PK brings DNA ends together, forming long-range, short-lived synapsis. In the presence of X-Ray Repair Cross Complementing 4 (XRCC4), XRCC4-like factor (XLF), DNA ligase 4 (Lig4) and with or without Paralog of XRCC4 and XLF (PAXX), a short-range, long-lived mature synapsis is achieved.^5,6^ Structural studies further show that DNA-PK dimerizes and bends the broken DNA ends at an angle of approximately 90 degrees^7^ and is present in the NHEJ supercomplex.^8,9^ However, an alternative NHEJ assembly is possible in the absence of DNA-PKcs.^10^ In addition, DNA-PKcs also initiates the DNA damage response by phosphorylating histone H2AX.^11,12,13^ Thus, we ask if DNA-PKcs functions as a hierarchical regulator that facilitates DSB repair in a timely manner to prevent DSB end uncoupling and illegitimate chromosome rearrangements.^14,15,16^

Inhibition of DNA-PKcs in mouse cells increases the likelihood of broken ends from different chromosomes to co-localize, which is thought to increase chromosome translocation.^17^ In the context of murine immunoglobulin heavy chain class-switch recombination, DNA-PKcs suppresses translocation to other genomic loci.^18,19^ However, the mechanisms of chromosomal translocation differ between mouse and human cells. In mice, alternative end joining (A-EJ) is the dominant repair machinery responsible for chromosomal translocation, characterized in cycling cells by increased microhomology-mediated end joining (MMEJ). ^20, 21^ In contrast, NHEJ is primarily responsible for chromosome translocation in human cells.^22^ It remains unknown how DNA-PKcs affects the chromosome translocation rate in human cells.

Unlike chromosome translocation, V(D)J recombination is a physiological and highly orchestrated chromosome rearrangement process that occurs within antigen receptor loci of developing B and T lymphocytes. This multi-step process begins with 1) acquisition of compatible recombination signal sequence (RSS) pairs regulated by cohesion complexes and chromatin loops;^23^ this is followed by 2) RAG1/2 (RAG)-mediated DSB generation at each RSS/gene segment interface, resulting in a pair of hairpin sealed coding ends and blunt signal ends, held in a postcleavage synaptic complex, where 3) NHEJ accesses and joins like ends to each other. Removal of the hairpin structure relies on the DNA-PKcs-dependent enzyme Artemis,^24^ highlighting the importance of DNA-PKcs in coding-coding joining. In mice, ablation of DNA-PKcs results in a massive failure of coding end joining, while signal end joining is partially affected,^25,26,27^ due to redundant functions with ATM.^28^ However, the comprehensive impact of DNA-PKcs depletion on coding-coding, signal-signal, and illegitimate chromosome rearrangements (i.e. coding-signal joints, translocations etc.) has not been fully evaluated.

Therefore, we systematically investigated DNA-PKcs functions in suppressing illegitimate chromosome rearrangements, and to what extent A-EJ contributes to that process by employing the <u>L</u>inear <u>A</u>mplification <u>M</u>ediated <u>H</u>igh-<u>T</u>hroughput repertoire, rejoin and <u>G</u>enome-wide <u>T</u>ranslocation <u>S</u>equencing (LAM-HTGTS) methodologies.^29–34^ In both human Cas9- and mouse RAG-mediated DSB cell lines, we find that incomplete chemical inhibition of DNA-PKcs leads to increased chromosome translocation of blunt ended DSBs that are repaired by NHEJ, whereas complete genetic inhibition suppresses NHEJ-mediated translocations and increases MMEJ-mediated translocations, manifested most strikingly in cycling cells. DNA-PKcs deficiency can still form NHEJ-mediated translocations of blunt ends, but hairpin sealed ends, in the context of V(D)J recombination, are transitioned into a phenotypic *Ku70^-/-^* A-EJ mechanism for repair. However, unlike *Ku70^-/-^*, DNA-PKcs deficiency does not restore the poor repair efficiency of *Lig4^-/-^*in non-cycling cells but does reduce their range of resected joints and converts their near-exclusive MMEJ activity into a more direct-dominant *Ku70^-/-^*junction structure pattern. These findings suggest that DNA-PKcs kinase activity is a crucial step to suppressing chromosome translocations and proceeding to complete repair by NHEJ that would otherwise be committed to repair involving MMEJ. They also suggest DNA-PKcs is not a major roadblock to A-EJ, but that it contributes to a Lig4-independent end joining pathway that engages DNA end resection and microhomology (MH) on protected ends with prolonged lack of repair.

## Methods

### Plasmids Preparation

The human Bcl2 gene (CDS) was cloned into a pLenti-puro vector to generate lentivirus. The Lenti-iCas9-Neo vector containing the Flag-iCas9-P2A-GFP cassette for Cas9 expression in K562 cells was obtained as a gift from Qin Yan (Addgene plasmid #85400,^35^). For Cas9 expression in HEK293T cells, the neomycin-resistant gene of Lenti-iCas9-Neo was replaced with a blasticidin-resistant gene, resulting in the Lenti-iCas9-Blast vector. The CRISPR-Cas9 Guide RNA expression vector pMCB320 (Addgene #89359,^36^) and the pMCB320-ACOC library (Addgene #101926,^37^) bearing a mCherry reporter were obtained as gifts from Michael Bassik. The pMCB320 vector was used to clone guide RNAs targeting RAG1D, RAG1L, and HBB sites, resulting in pMCB320-RAG1D, pMCB320-RAG1L, and pMCB320-HBB, respectively.

### Cell Lines Modification

The human immortalized myelogenous leukemia cell line, K562, derived from a CML patient, (ATCC #CCL-243) was used in this study. K562-*Bcl2* cells were generated by transducing K562 cells with lentivirus bearing human Bcl2. K562-*PKcs^-/-^* cells were generated by introducing a DNA-PKcs exon-77 deletion using CRISPR-Cas9 in K562-*Bcl2* cells and verified by PCR, sanger sequencing and western blot (see the main text).

K562-*iCas9* cells were generated by transducing K562 cells with lentivirus bearing iCas9-Neo. The K562 cell line and its modified versions were cultured at 37°C and 5% CO2 in RPMI-1640 medium supplemented with 10% (vol/vol) fetal bovine serum (FBS), 50 U/mL penicillin/streptomycin/2 mM L-glutamine (Gemini Bio-Products, # 400-110), 1× MEM-NEAA (Lonza, #13-114E), 1 mM sodium pyruvate (Cytiva, #SH30239.01), 50 μM 2-mercaptoethanol (Sigma-Aldrich, #M6250-100ML), and 20 mM HEPES (pH 7.4, Cytiva, #SH30237.01). K562-*Bcl2* and K562-*PKcs^-/-^* cells were selected with 5 μg/ml puromycin (Sigma-Aldrich, #P8833-100MG), and K562-*iCas9* cells were selected with 200 μg/ml G418 (BioVision, #2864-20). Doxycycline (2 μg/ml, Sigma-Aldrich, #D9891-10G) was added only when Cas9 expression was required. Doxycycline-treated cells were not used for cell line preservation.

The human embryonic kidney cell line, HEK293T, is an SV40 Large T antigen transformed derivative HEK293 epithelial cell line (ATCC #CRL-3216) was used in this study. HEK293T-*iCas9* cells were generated by transducing HEK293T cells with lentivirus containing iCas9-Blast. HEK293T and HEK293T-*iCas9* cells were cultured at 37°C and 5% CO2 in DMEM supplemented with 10% (vol/vol) FCS, 50 U/mL penicillin/streptomycin, 2 mM L-glutamine, 1× MEM-NEAA, 1 mM sodium pyruvate, 50 μM 2-mercaptoethanol, and 20 mM HEPES (pH 7.4). HEK293T-iCas9 cells were selected with 10 μg/ml blasticidin. Doxycycline (2 μg/ml) was added only when Cas9 expression was required. Doxycycline-treated cells were not used for cell line preservation.

The human colorectal carcinoma cell line, HCT116, with various DNA-PKcs gene editing including DNA-PKcs^+/-^, DNA-PKcs^-/-^ and DNA-PKcs^KD/-^ were generated and verified as previously described.^38^ The DNA-PKcs-modified HCT116 cell lines were cultured at 37°C and 5% CO2 in Hyclone α-minimum Eagle’s medium supplemented with 5% newborn calf serum, 5% fetal bovine serum (Sigma), and 1× penicillin/streptomycin (Gibco).

Mouse Abelson kinase-transformed progenitor B cell (*v-Abl)* lines including wild-type (*WT*), *Ligase 4^-/-^* (*Lig4^-/-^*), *Ku70^-/-^*, and *Lig4^-/-^Ku70^-/-^* cells were reported in our previous study (clone A).^29^ To generate new cell lines, DNA-PKcs was deleted using CRISPR/Cas9 in *WT* and *Lig4^-/-^* cells, resulting in cell lines *DNA-PKcs^-/-^* (#1/2) and *Lig4^-/-^PKcs^-/-^*(#1/2), respectively. Confirmation of these cell lines was performed by genotyping using the primers listed in Table S7. All *v-Abl* cells were cultured at 37°C and 5% CO_2_ in RPMI-1640 medium supplemented with 10% (vol/vol) FBS, 50 U/mL penicillin/streptomycin/2 mM L-glutamine, 1× MEM-NEAA, 1 mM sodium pyruvate, 50 μM 2-mercaptoethanol, and 20 mM HEPES (pH 7.4). To induce V(D)J recombination, *v-Abl* cells were seeded at a concentration of 1 million cells/ml and supplemented with 3 μM STI-571 (TCI Chemicals, #I0936-100MG) for four days. Cells treated with STI-571 were not used for cell line preservation.

Mouse DNA-PKcs catalytically dead *v-Abl* cell lines, *DNA-PKcs^KD/KD^* #C1, #C3 and #C3-1 were generated and verified as previously reported.^39^ The cell culture conditions were the same as above for *v-Abl* cells except the base media DMEM.

### Lentivirus Production

Lentivirus was generated using the 2nd generation lentivirus packaging system. The transfer plasmid containing the gene or guide RNA of interest, the packaging plasmid psPAX2, and the envelope-expressing plasmid pMD2.G were mixed at a 4:3:1 mass ratio, which corresponded to 2 μg, 1.5 μg, and 0.5 μg, respectively, in 120 μl of opti-MEM (Fisher Scientific #31-985-070). To this mixture, 5 μl of the p3000 reagent was added and mixed thoroughly. In a separate tube, 5 μl of lipofectamine 3000 (Thermo Scientific, #L3000015) was mixed with 120 μl of opti-MEM and mixed thoroughly. The DNA mixture was then added drop by drop to the lipofectamine mixture, mixed well, and left at room temperature for 15 mins.

Next, the lipofectamine-DNA mixture was added to one well of HEK293T cells cultured in a 6-well plate and maintained in culture media for an additional two days. The media containing lentivirus was collected on day 2, and 2.5 ml of fresh media was added to the cells, which were maintained for one more day. The media was then collected again to obtain more lentivirus. The lentivirus titration was recommended to be performed in the targeting cell lines rather than in HEK293T cells alone.

### Immunoblotting

The samples, including K562-*WT*, K562-*Bcl2*, K562-*PKcs^-/-^*, and K562-*iCas9*, were collected at a concentration of 5 million cells and resuspended in 200 μl of RS buffer (150 mM NaCl, 10 mM Tris pH 7.5). Cell lysis was performed by adding 200 μl of 2X Laemmli buffer (4% SDS, 5% 2-mercaptoethanol, 20% glycerol, 0.004% bromophenol blue, 125 mM Tris pH 6.8) and incubating at 95°C for 10 minutes. The lysate samples were loaded directly onto a 4-15% TGX precast SDS-PAGE gel (Bio-Rad, #4561084). Protein transfer onto a nitrocellulose membrane was carried out using the Turbo™ Transfer System (Bio-Rad, #1704150) and the corresponding transfer kit (Bio-Rad, #1704271). Immunoblotting was performed using the following antibodies: anti-Bcl2 (1:2000, Novus Biologicals #NB100-56098), anti-Flag (1:2000, Cell Signaling Technology #12793S), anti-DNA-PK (1:4000, Origene #TA314389), anti-Rabbit-IgG (1:2000, Thermo Scientific #G-21234), and anti-actin (1:4000, Santa Cruz Biotechnology #SC-47778). Protein band detection was accomplished using an HRP substrate (Advansta #K-12042-D10) and visualized under a chemiluminescent imaging system, ChemiDoc (See Figures S1A, S1D). The Cas9 expression level in HEK293T-iCas9 cells was detected using the same approach (See Figure S1G). The phosphorylation state of DNA-PKcs S2056 (PKcs p^S2056^) was detected using the corresponding antibody (Thermo Scientific #PA5-121294) in similar manner, except phosphatase inhibitors were added in cell lysis. After detection of PKcs p^S2056^, its antibody was removed by stripping buffer (Takara Bio #T7135A) and re-blotted by PKcs antibody to quantify the total amount of DNA-PKcs (See Figures S2I-L).

### Quality Control by Flow Cytometry

#### Cell Cycle Analysis

The cell cycles of K562-*Bcl2* cells, with or without Palbociclib, DPKi #1/2, and ATMi, were assessed based on DNA replication activity. Briefly, 50 μM EdU (Cayman #20518) was added to each condition and incubated for 30 minutes. The EdU-treated cells were fixed with 2% formaldehyde (EMS #15710) for 15 minutes at room temperature, permeabilized with 0.5% TritonX-100 for 15 minutes at room temperature and subjected to click labeling reaction using AFDye 488 Azide (Click Chemistry Tools, #1314). The AFDye 488-labeled cells were analyzed by flow cytometry using the FITC channel.

#### Plasmid Delivery into K562 and HCT116 cell lines

For delivering plasmids into K562-*Bcl2* and K562-*iCas9* cells, the 4D-nucleofector system (Lonza, Core plus X unit) and SF Cell Line 4D X Kit (Lonza, #V4XC-2024) were employed. Four million K562-*Bcl2* cells were nucleofected with 5 μg pX330-RAG1D, 5 μg pX330-RAG1L and 0.5 μg pMAX-GFP. The cells were then cultured for two days. Nucleofection efficiency was determined by measuring GFP expression from the pMAX-GFP vector using the FITC channel of flow cytometry. Similarly, four million K562-*iCas9* cells were nucleofected with 5 μg pMCB320-ACOC plus 5 μg pMCB320-RAG1L or 5 μg pMCB320-ACOC plus 5 μg pMCB320-HBB. The cells were cultured for two days, and nucleofection efficiency was evaluated by detecting mCherry expression from the pMCB320 vector using the Cy5 laser channel of flow cytometry. The expression level of Cas9 upon Doxycycline induction was correlated with GFP expression detected by the FITC channel. Plasmids including pX330-RAG1D, pX330-RAG1L and pMAX-GFP were delivered into HCT116 cells same as K562-*Bcl2* cells but using the SE Cell Line 4D-Nucleofector™ X Kit (V4XR-1024), and nucleofection efficiency was evaluated similarly as above.

#### Plasmid Transfection into HEK293T-*iCas9* Cells

HEK293T-*iCas9* cells were transduced in 6-well plates with 2.5 μg pMCB320-RAG1D and 2.5 μg pMCB320-RAG1L or with 4 μg pMCB320-ACOC and 1 μg pMCB320-RAG1L, using the lipofectamine 3000 kit. The cells were maintained in D10 media, with or without DPKi #1/2. Transfection efficiency was determined by measuring mCherry expression from the pMCB320 vector using the Cy5 laser channel of flow cytometry. The expression level of Cas9 upon Doxycycline induction was correlated with GFP expression detected by the FITC channel.

### LAM-HTGTS Library Preparation

HTGTS library preparation was performed as previously described^31^ with some modifications. For K562-*Bcl2*, K562-*iCas9*, HEK293T-*iCas9* and HCT116 libraries, 5 μg of genomic DNA from each treatment condition was sheared using a bioruptor sonication device (Diagenode) in low mode for two cycles (30s on + 60s off) at 4°C, resulting in fragments ranging from 200 bp to 2 kb. The sheared fragments were subjected to linear amplification (LAM)-PCR using biotin-labeled primers, including Bio-RAG1L and Bio-HBB proximal to the RAG1L and HBB cleaved sites (baits), respectively. The LAM-PCR products were enriched using streptavidin-coated magnetic beads, followed by adapter ligation. Unligated adapters were removed, and the ligated products were subjected to nested-PCR using a common primer (AP2I7-novo) matching the adapter sequence and another barcoded I5 primer that matches the region between the bait-site and the biotin-labeled primer. The DNA from the nested-PCR was purified using a Gel-extraction kit (Qiagen #28706). Subsequently, tagged PCR was performed using primers P7I7 and P5I5, which match the primers used in the nested-PCR. The PCR products were purified by 1% agarose gel electrophoresis, and DNA products with a length of 500 bp to 1 kb were excised and extracted using a Gel-extraction kit. The tagged DNA libraries were subjected to bioanalyzer analysis for quality control and sequencing using the Illumina NovaSeq-PE150. Please refer to Table S6 for the oligos used.

The HTGTS library of mouse *v-Abl* cells was prepared using a similar approach, with the following modifications: 1) 2 μg of genomic DNA/sample was used for each library; 2) Barcoded AP2I7-novo primers were used, and the nested-PCR products from each of the 12 libraries were pooled and tagged ensemble in tagged PCR; 3) Jκ1, Jκ2, Jκ4, and Jκ5 coding and signal end-specific biotin-labeled primers, as well as nested I5 primers, were used. Please refer to Table S6 for the oligos used.

### Data Analyses

LAM-HTGTS data analysis was performed following previously reported methods.^29,31^ Briefly, sequencing reads from Illumina NovaSeq PE150 were de-multiplexed based on the inner barcodes and the sequence between the bait site and the nested PCR I5 primers using the fastq-multx tool from ea-utils. The adapter sequences were trimmed using the SeqPrep utility. The demultiplexing and trimming functions were integrated into a script called TranslocPreprocess.pl. Subsequently, the reads were normalized using Seqtk and mapped to the hg19 or mm9 reference genome using TranslocWrapper.pl to identify chromosome translocations or V-J recombination events, generating result tlx files. Junctions that aligned to the bait region were not shown in the result tlx files and were extracted separately using a script called JoinT.R, resulting in final tlx files containing translocations and rejoin events. JoinT.R was omitted for V-J recombination analysis. Junction duplications were retained as described previously.^29,33^

Result and final tlx files were converted into bedgraph files using tlx2bed.py,^31^ which were then visualized and plotted using IGV (integrative genomics viewer). Junctions in regions of interest from the result and final tlx files were extracted using tlxbedintersect.py, which relied on two other scripts, tlx2BED.pl and pullTLXFromBED.pl.^31^ The regions of interest varied depending on the specific paradigm. For Vκ-Jκ recombination in *v-Abl* cells, the regions of interest were the RAG1/2 cleavage sites of Vκ genes, with a flanking 200 bp window (± 200). In the case of twinned Cas9-cleaved K562-*Bcl2*, the region of interest was the RAG1D “prey” locus. In the genome-wide DSBs generated by ACOC libraries in K562-*iCas9* and HEK293T-*iCas9* cells, the final tlx files were converted into bed files using tlx2BED-MACS.pl and evaluated using the peak caller MACS2, resulting in a peaks.xls file that contained the junction peaks and their significance. The peak regions in control and experimental conditions were combined together and used as regions of interest.

Junctions were plotted using TranslocPlot.R, ^14^ which provided visualization of the chromosomes and the locations of junctions across the genome. These commands could be used for plotting junctions across the entire genome or within a specific region on a given chromosome. Circos plots were generated using the circos package, combining the binfile.txt from the junction plot and the count.bed from tlxbedintersect.py. The result and final tlx files could also be utilized for repair pathway and resection analyses. JctStructure.R was employed to determine the repair patterns, including microhomology, direct repair, and insertion. The degree of DSB end resection, indicated by the distribution of junctions near the DSB break site, was quantified using ResectionRSS.R.

The data obtained from western blot and flow cytometry experiments were analyzed using ImageJ (NIH) and FlowJo (FlowJo, LLC), respectively.

### Statistical Analysis

All data were presented as mean ± SEM if not otherwise indicated. Analyses of differences were determined using one-way ANOVA plus Dunnett’s multiple comparisons, or Student’s t-test (unpaired Student’s t-test for the data obtained in same batch and ratio paired Student’s t-test for the data obtained in different batches). Statistical analyses were performed using GraphPad Prism (GraphPad Software, Inc.). Where necessary, linked experimental sets were displayed to discern batch effects. All P values presented were two-tailed, and P < 0.05 was considered statistically significant.

## Results

The significance of DNA-PKcs in physiological DNA damage repair has led to the development of inhibitors for cancer treatment.^40–44^ For this study, we chose two well-established inhibitors, Nu7441 (DPKi #1 at 500nM) and Nu7026 (DPKi #2 at 20μM). As ATM has been extensively characterized in suppressing chromosome translocation,^45^ we also included Ku-60019, a specific ATM inhibitor (ATMi at 10μM), as a control.^46^ Inhibitor concentrations were determined by the minimal amount that promoted chromosome translocations in pilot experiments (data not shown). We used the CRISPR-Cas9 system in K562 and HEK293T cell lines (Figures S1A-S1I) (see methods). To minimize cell death by nucleofection, we introduced the anti-apoptosis gene Bcl2 into K562 cells, resulting in the generation of a novel cell line, K562-*Bcl2*. Western blot analysis confirmed significantly higher expression levels of Bcl2 in K562-*Bcl2* cells compared to the parental cell line, K562-*WT* (Figure S1A), and the efficiency of nucleofection was assessed by co-nucleofecting a GFP vector along with CRISPR-Cas9 (Figures S1B, S1C). We separately generated inducible Cas9-expressing cell lines, K562-*iCas9* and HEK293T-*iCas9*, and confirmed Cas9 expression upon induction with Doxycycline using western blot analysis (Figures S1D, S1G). Guide RNA (gRNA) pools were delivered^37^ with high level gRNA expression (Figures S1E-S1F, S1H-S1I).

We first confirmed the effect of these inhibitors did not drastically affect cell cycle progression (Figures S2A-S2D) and then next investigated whether the treatment with these inhibitors would have a similar affect under G1/G0 arrest using the cdk4/6 inhibitor, Palbociclib.^47,48^ G1/G0 arrest was achieved with 50μM Palbociclib in K562 cells, and the inclusion of DPKi #1, DPKi #2, or ATMi did not interfere with the cell cycle arrest induced by Palbociclib (Figures S2E-S2H). To confirm the efficacy of the DNA-PKcs inhibitors, we performed immunoblotting to detect the total level of DNA-PKcs and its auto-phosphorylated form at Ser2056 (pS2056). We observed a marginal decrease in total DNA-PKcs levels, while the DNA-PKcs auto-phosphorylation at Ser2056 was significantly reduced upon treatment with DPKi #1 and DPKi #2 in cycling K562-*Bcl2* cells (Figures S2I-S2J), as well as in Palbociclib-treated cells (Figures S2K-S2L). Based on these results, we conclude that DPKi #1 and DPKi #2 do not perturb the cell cycle in the context of our study.

### Increased Intrachromosomal Translocation by Partial DNA-PKcs Inhibition

Using nucleofection in K562-*Bcl2* cells, we employed a pair of Cas9:gRNAs in the RAG1 locus (RAG1D and RAG1L) on chromosome 11 to generate DSBs using the RAG1L cleavage site as the bait, enabling the detection of junctions arising from single (rejoined) or dual (translocated to RAG1D) Cas9-generated DSBs via HTGTS-JoinT-seq^29^ (Figure 1A), which for the latter would include deletions and inversions based on chromosome orientation. Repair patterns of these DSBs are consistent with the generation of predominantly blunt ends.^49^ For these experiments, we derived a translocation rate (TL) by normalizing against the number of rejoin events (Figure 1A). In cycling K562-*Bcl2* cells, we generated hundreds of thousands of junctions (Tables S1 and S2) and observed an increased number of translocations in the presence of DNA-PKcs inhibitors, DPKi #1 and DPKi #2, compared to the DMSO control that was accompanied by a decrease in the number of rejoins (Figure 1A). Conversely, ATMi treatment resulted in a slight decrease in the number of translocations but with a significant drop in rejoins that was on average 50% greater than with DPKi treatments (Figure 1A). Consequently, all inhibitors led to an elevated translocation rate, with a two-fold increase in ATMi and DPKi #1, and a three-fold increase in DPKi #2, compared to the DMSO control (Figure 1B). Previous work demonstrated that non-homologous end joining (NHEJ) is responsible for chromosome translocation in human cells.^22^ To investigate if the observed increase in chromosome translocation from inhibited DNA damage signaling was due to A-EJ, we analyzed junction structures of the resulting translocations. Unexpectedly, joints remained predominately direct (Figure 1C), suggestive of continued utilization of NHEJ.

**Figure 1.**
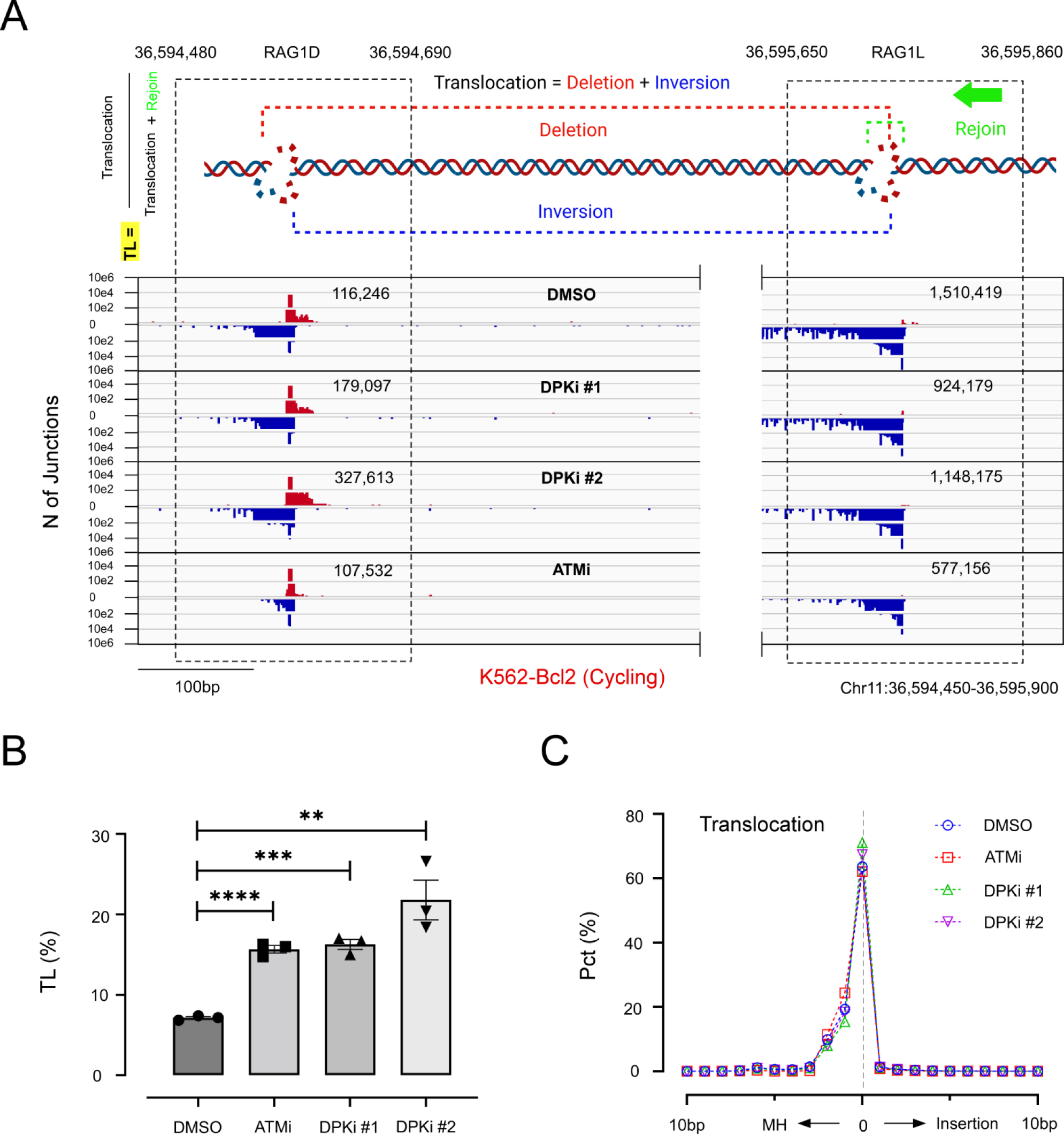
Increased NHEJ-Mediated Intrachromosomal Translocations by Partial DNA-PKcs Inhibition in Cycling K562-Bcl2 Cells. (**A**) Cas9:RAG1D “prey” and Cas9:RAG1L “bait” DSBs on chromosome 11 for high-throughput genome-wide translocation sequencing (HTGTS; green arrow). Three dominant types of repair junctions were formed: deletion (red dashed line), inversion (blue dashed line) and rejoin (green dashed line). Cas9:RAG1D translocation rate (TL): (deletions + inversions) / (rejoin + deletions + inversions). IGV plots display junction profiles of the RAG1D and RAG1L sites for DMSO, DPKi #1, DPKi #2 or ATMi conditions. (**B**) TL rates for the tested conditions: DMSO (7.15 ± 0.16%), ATMi (15.68 ± 0.47%), DPKi #1 (16.26 ± 0.62%), and DPKi #2 (21.79 ± 2.46%). Statistical significance between DMSO and inhibitors was assessed using unpaired T-test; **p<0.01; ***p<0.001; ****p<0.0001. (**C**) Percent Translocation junction structure distributions were categorized into bps of microhomology (MH), insertion, and direct repair (“0”). Experiments were from three biological replications, and error bars using standard error of the mean (SEM).

We next investigated whether DNA-PKcs inhibition affected chromosome translocation in G1/G0-arrested K562-*Bcl2* cells. Under these conditions we accumulated ∼2-fold fewer total junctions compared to the cycling counterparts (Tables S1 and S2). Similar to our cycling experiments (Figures 1A-1C), DPKi #1 and DPKi #2 increased the number of translocations and reduced the number of rejoins compared to the DMSO control, while ATMi decreased both the number of translocations and rejoins (Figure 2A). Consequently, the translocation rate was significantly higher for DPKi #1 and DPKi #2 while ATMi showed a slight upward trend compared to the DMSO control (Figure 2B). Further analysis of translocation junction structures did not reveal any changes relative to the control, which were direct and indicative of NHEJ (Figure 2C).

**Figure 2.**
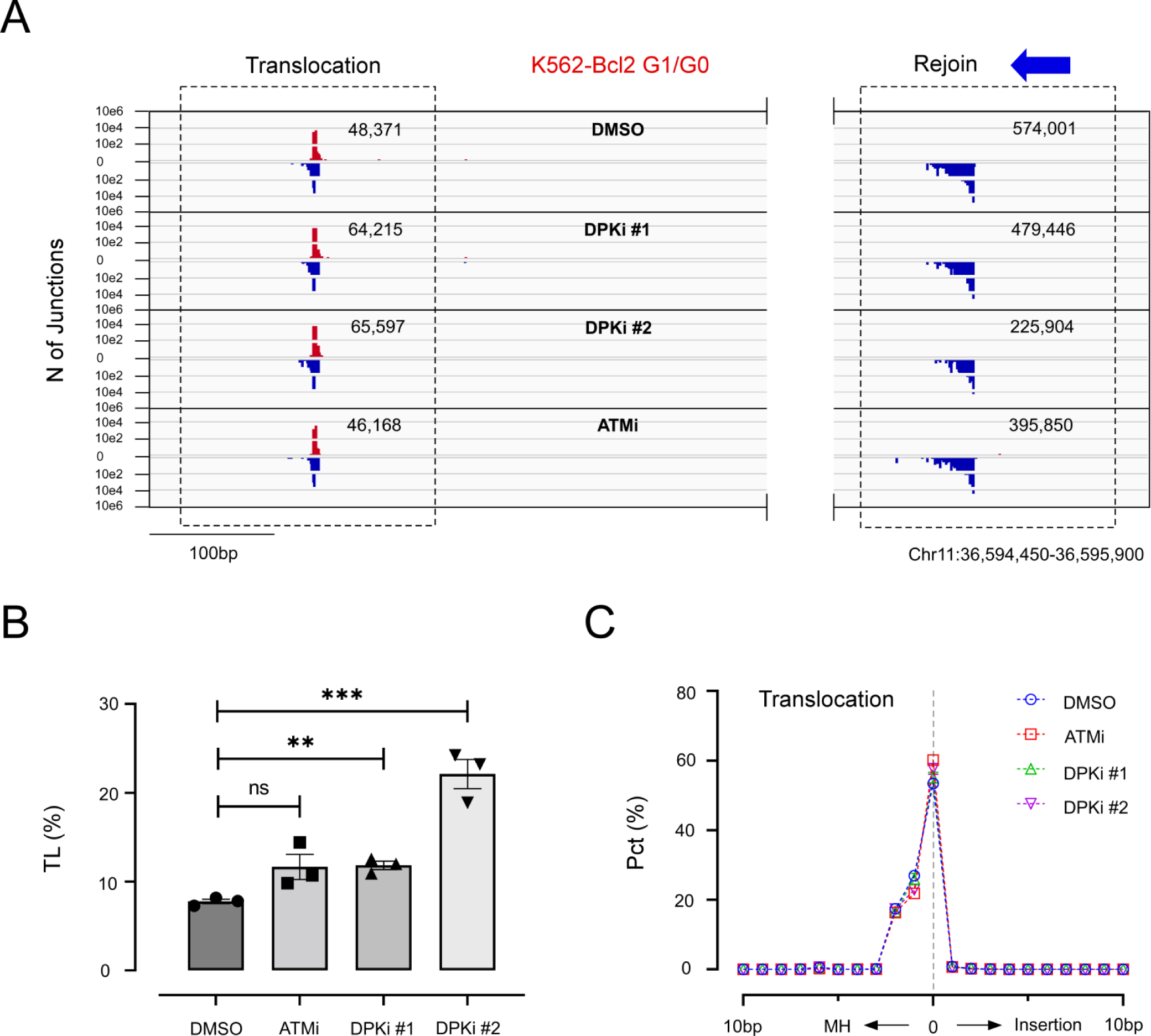
Partial DNA-PKcs Inhibition Promotes NHEJ-Mediated Intrachromosomal Translocations in G1/G0-Arrested K562-Bcl2 Cells. **(A)** IGV junction profile plots of RAG1D and RAG1L sites in the presence of cdk4/6 inhibition (Palbociclib) and treated with DMSO, DPKi #1, DPKi #2, or ATMi. Blue arrow indicates the bait site. **(B)** TL rates for DMSO (7.78 ± 0.26%), ATMi (11.66 ± 1.41%), DPKi #1 (11.85 ± 0.47%), and DPKi #2 (22.12 ± 1.64%), evaluated by unpaired T-test; **p<0.01; ***p<0.001; ns, not significant. **(C)** Translocation junction structure distributions; see Figure1 legend for details. Experiments were independently repeated three times, and SEM are provided.

From the above findings, chromosome translocations are mediated by NHEJ, regardless of the presence or absence of DNA-PKcs inhibitors; however, Ku70 deficiency in G1/G0-arrested *v-Ab*l cells,^29^ representing a robust A-EJ pathway that also leverages direct joining, may not be readily distinguished if DPKi effects favor this pathway over NHEJ. To test for this possibility, we inhibited two crucial components of A-EJ, poly(ADP-ribose) polymerase-1 (Parp1)^50,51^, and polymerase theta (Polθ),^52^ using Olaparib (Parpi at 10μM)^53^ and Novobiocin (Polθi #1 at 100μM)^54^ in G1/G0-arrested K562-*Bcl2* cells, with or without the DNA-PKcs inhibitor DPKi #2. Here, DPKi #2 had the most pronounced effect, leading to an increase in translocations and a decrease in rejoins (Figure S3A). Parpi and Polθi #1 slightly elevated both translocations and rejoins, irrespective of the presence or absence of DPKi #2. Consequently, we observed no significant change in the translocation rate with Parpi or Polθi #1 alone or when combined with DPKi #2 (Figure S3B). Further analysis of the Cas9 translocation repair pattern did not reveal any significant changes (Figure S3C).

To validate our findings in a different cell line, we performed similar experiments in HEK293T-*iCas9* cells, where the same pair of gRNAs were used (RAG1D and RAG1L) for targeted DSB generation upon doxycycline induction (see methods). We observed a similar increase in translocation events upon DNA-PKcs inhibition, with no alterations in translocation junction structures (Figures S4A-S4C), and similar effects were observed upon ATMi treatment (Figure S4D, S4E) (Tables S1 and S3). Thus, we conclude that DNA-PKcs inhibition at the doses assayed promotes proximal chromosome translocation by NHEJ.

### Partial DNA-PKcs Inhibition Promotes Genome-wide Translocations

We investigated whether the phenomenon observed with the Cas9:RAG1D prey site applies more broadly, by generating genome-wide double-strand breaks (DSBs) using a gRNA library, ACOC, which contains approximately 30,000 different gRNAs.^37^ These DSBs were introduced in G1/G0-arrested K562-*iCas9* cells while using the Cas9:RAG1L DSB as bait for JoinT-seq library preparation and recovered ∼200,000-300,000 total junctions per replication (Tables S1 and S3). Considering recent findings highlighting the role of topologically associated domains (TADs) in DNA damage repair,^55^ we defined TAD boundaries based on CTCF and RAD21 chip-seq data (Figure S5A) and evaluated junctions located beyond the RAG1 TAD. Quantification of these genome-wide translocations and bait DSB rejoins clearly indicate significantly more translocations and less rejoins for DPKi #1 and DPKi #2 conditions relative to DMSO control (Figure 3A). Applying a similar TL measure but now for translocations exterior to the RAG1 TAD, exTAD-TL, was >2-fold higher in the DPKi #1 and DPKi #2 conditions compared to the DMSO control (Figure 3B). Importantly, this more diverse collection of Cas9:gRNA-targeted chromosome translocations were still predominantly direct, indicative of NHEJ, and with no significant perturbation observed upon DPKi #1/2 treatment (Figure 3C). Genome-wide translocation analysis was also conducted in cycling HEK293T-*iCas9* cells, and consistent results were obtained (Figure S6A; Tables S1 and S3), including no significant changes in junction structure patterns (Figure S6B-S6C).

**Figure 3.**
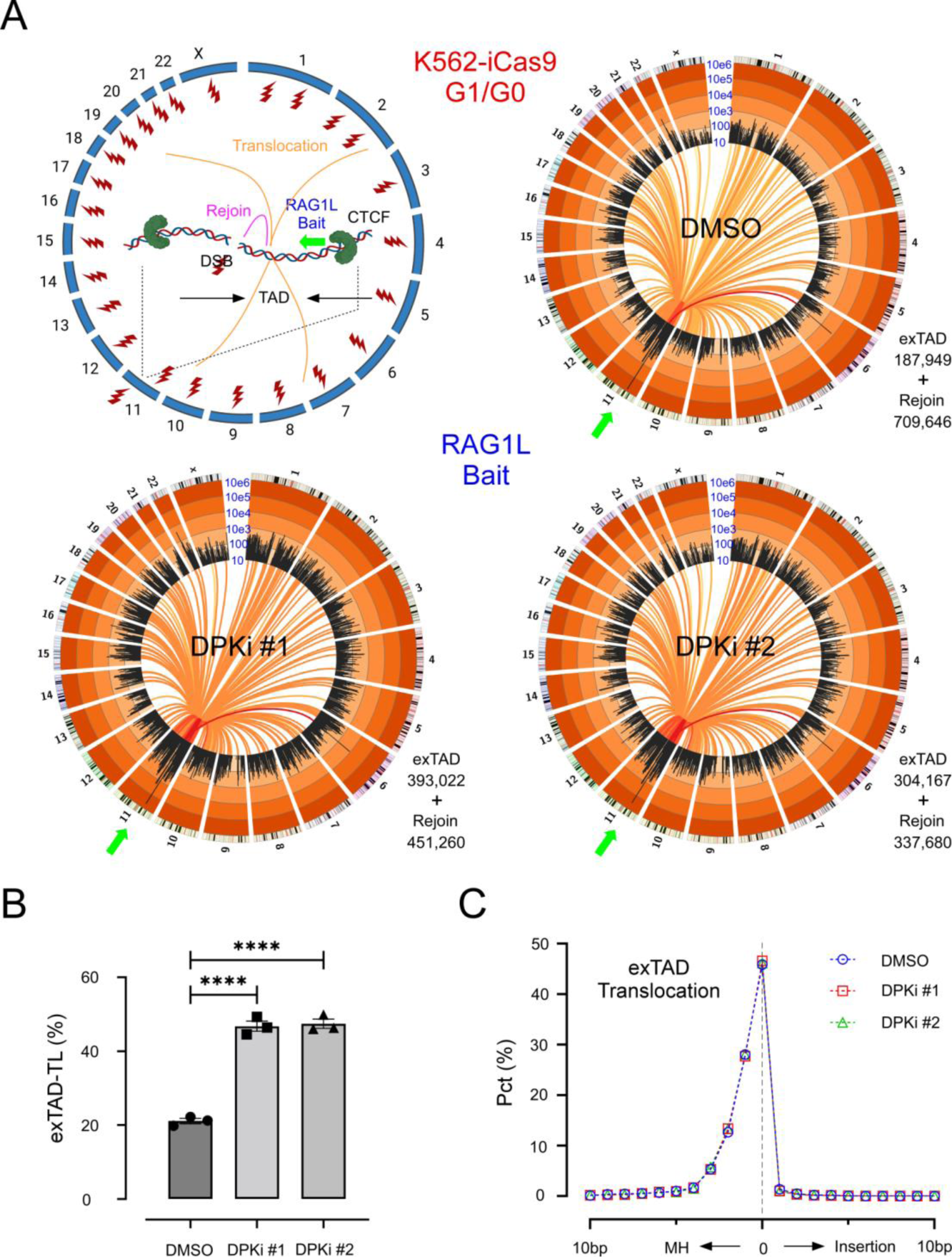
Partially inhibited DNA-PKcs Promotes Genome-wide Translocations. (**A**) To generate double-strand breaks (DSBs), ∼30,000 ACOC library gRNAs (represented as red lightning bolts) were transfected in G1/G0-arrested *K562-iCas9* cells with doxycycline induction of Cas9. The RAG1L gRNA was used as bait for HTGTS (green arrows). (Top left) Translocations and rejoins are depicted by orange and magenta lines, respectively. The bait site is flanked by convergently oriented CTCF binding sites, forming a Topology-Associated Domain (TAD, Chr11: 36,585,950-36,644,640). The genome-wide translocation profiles for DMSO, DPKi #1, and DPKi #2 conditions are shown as circos plots with junction frequencies reported within (Rejoin) or genome-wide (exTAD). Black bars indicate 5Mb binned chromosome regions and their frequency is indicated on log scale. Yellow to red colored lines connect bait to prey hotspots with red being the most significant. (**B**) The TL rates measuring junctions outside the TAD region (exTAD) normalized to the number of rejoins within the TAD (exTAD-TL). The exTAD-TL changes between DMSO (21.10 ± 0.71%), DPKi #1 (46.76 ± 1.38%), and DPKi #2 (47.43 ± 1.24%) were evaluated by unpaired T-test. ****p< 0.0001. (**C**) Junction structure distributions for exTAD translocations; see Figure 1 legend for more details. Experiments were independently repeated three times, and SEM are provided.

To determine if the increased translocation upon DNA-PKcs inhibition was specific to the previous bait site, we employed a separate bait site in the HBB locus on chromosome 11 which was then subjected to the same ACOC gRNA library and assayed in G1/G0 arrested K562-*iCas9* cells (Figure S7A; Tables S1 and S3). The TAD domain encompassing the HBB locus was determined based on CTCF binding sites (Figure S5B). Again, significantly increased exTAD-TL values were observed in DPKi #1 and DPKi #2 (Figure S7B), while no changes were found in the repair patterns (Figure S7C). These findings suggest the increased chromosome translocation induced by DNA-PKcs inhibitors is a general phenomenon.

### Complete DNA-PKcs Inhibition Promotes MMEJ-Mediated Translocations

Although DPKi #1/2 inhibitors enhanced the translocation effect with no discernable change in junction structure patterns, autophosphorylation levels were not completely absent (Figures S2I-S2J), indicating incomplete kinase inhibition. Thus, we retested DNA-PKcs inhibition with a 10-fold increased concentration of DPKi #1 (5µM) and included two more potent DNA-PKcs inhibitors, DPKi #3 (AZD7648, 10µM) ^56,57^ and DPKi #4 (M3814, 10µM)^56,58^; in parallel to this experiment, we also tested the effect of a second Polθ inhibitor, ART558 (Polθi #2 at 10μM),^59^ in combination with DPKi #2. In G1/G0 arrested K562-Bcl2 cells, Polθi #2 displayed no significant change in translocations alone or with DPKi #2 and did not alter junction structure patterns (Figure 4A-4C). The higher dose of DPKi#1 and the effect of DPKi #3 and #4 increased TL rates, as expected (Figure 4A), however, the dosing efficacy of DPKi #1, 3 & 4 also significantly decreased translocation frequency (Figure 4B), indicating a substantial block in NHEJ. It is notable that no obvious cell death was detected within two-day experiment window. Junction structure patterns from interchromosomal translocations revealed a slight increase in MMEJ for DPKi #3/4 conditions (Figure 4C), which were the most affected. Importantly, MMEJ utilization became strikingly near-exclusive when DPKi #3/4 were tested again under cycling conditions despite having significant reductions in translocation frequency (Figure 4D-4F). We conclude a more robust chemical inhibition of DNA-PKcs kinase activity suppresses NHEJ-mediated repair and will ultimately transition to using MMEJ, particularly in cycling conditions.

**Figure 4.**
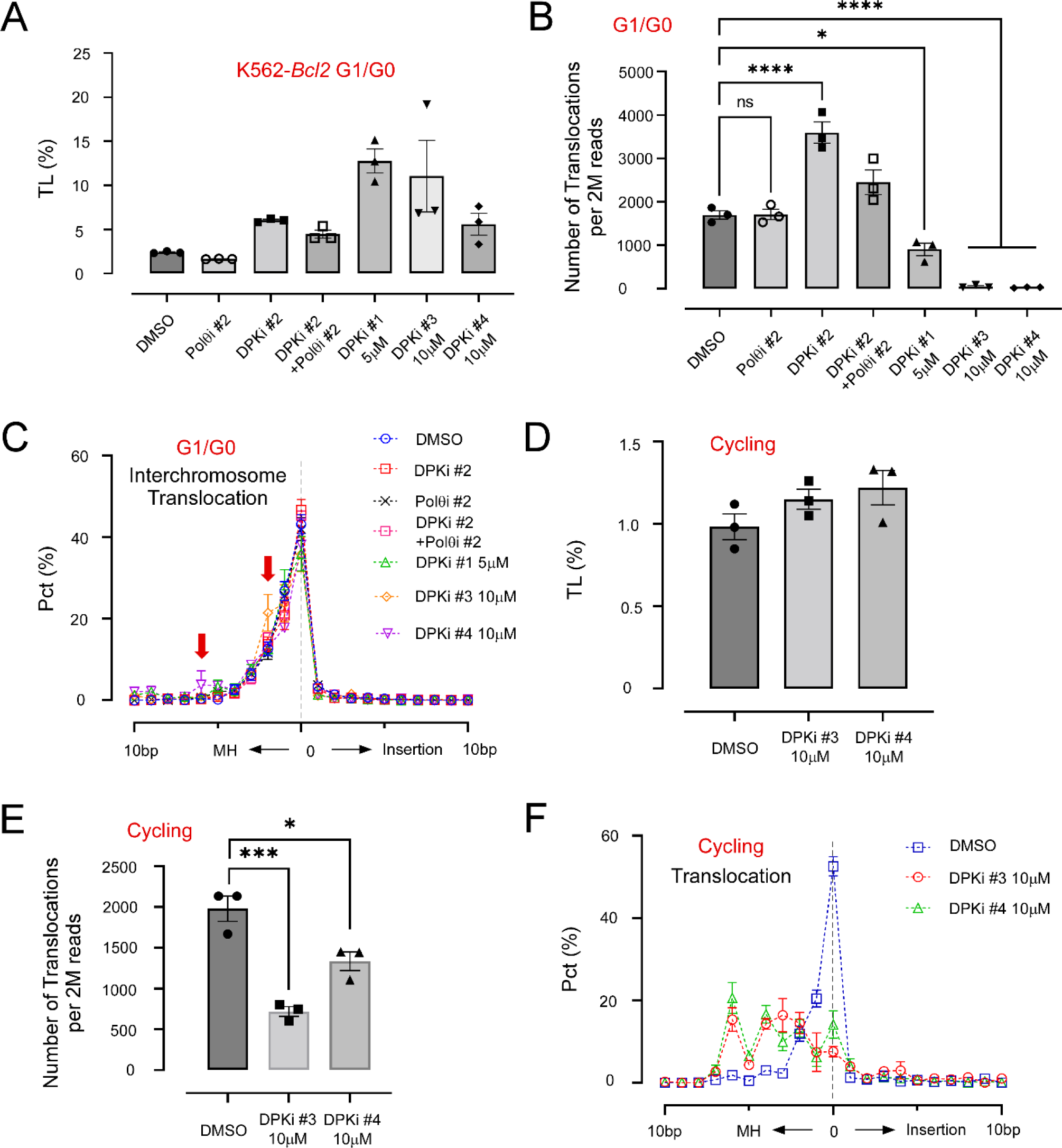
Complete DNA-PKcs Inhibition Promotes MMEJ-Mediated Translocations. (**A**) The TL rates between RAG1D and RAG1L sites of G1/G0-arrested *K562-Bcl2* cells upon treatment of DMSO, DNA-PKcs inhibitors DPKi #1-4, Polθi #2 and Polθi #2 + DPKi #2. Note, DPKi #1 was 5 µM, higher than the typical 0.5 µM for default assays. (**B**) The raw number of translocations out of 2 million total reads were shown for each condition and the difference was evaluated by one-way ANOVA plus post-comparison. (**C**) The junction structure in each condition with red arrow indicated the increased microhomology usage. (**D-F**) The two most potent DNA-PKcs inhibitors DPKi #3/4 were further tested in *K562-Bcl2* cycling cells, with (**D**), (**E**) and (**F**) same as (**A**), (**B**) and (**C**), respectively, but for the cycling. All the experiments were independently repeated three times (n=3), and the standard error of the mean (SEM) is indicated, with *p<0.05, ***p<0.001, ****p<0.0001 and ns (no significance).

We next wanted to contrast repair outcomes with DNA-PKcs deficiency by generating in K562-*Bcl2* cells a deletion in the kinase domain of DNA-PKcs (Δexon-77). Western Analysis indicated the deletion generated a truncation at a markedly low level (Figure S8A, S8B). While recovered total junctions with kinase domain deletion in G1/G0-phase were robust and averaged ∼83,000-108,000 (Figure S8C), the translocation rate and junction structure patterns remained unchanged with DPKi #1 (0.5µM) or DPKi #2 treatments (Figure S8D-S8E). However, DNA-PKcs deficiency revealed increased MMEJ from the Cas9:RAG1D prey DSB, although the major peak still corresponded with direct repair; extended analysis to interchromosomal translocations also revealed a doubled increase in MH utilization (MH > 3bp) in the PKcs^-/-^ versus the parental line (Figure S9A-S9B). Consequentially, we found Parpi, but not Polθi #2, reduced MMEJ and increased direct repair despite neither inhibitor affecting the TL rate (Figure S8F-S8G). Therefore, we conclude hypomorphic kinase truncation of DNA-PK promotes some MMEJ that involves Parp activity.

A prior report was not able to derive clonal lines of early exon DNA-PKcs deletions in HCT116 cells,^60^ which was analogous to our efforts in K562 cells but was able to do so with late exon deletions. More recently, sets of full DNA-PKcs deletion (in or after exons coding for the C-terminal kinase domain) and DNA-PKcs catalytically dead point mutations were reported.^38,61^ We applied our dual Cas9:gRNA approach to cycling HCT116 *DNA-PKcs^+/-^*, *DNA-PKcs^-/-^* and *DNA-PKcs^KD/-^* lines ^38^,which harbor an exon-79 deletion or K3753R kinase dead (KD) point mutation in exon-79, to measure changes in repair outcomes. Western Analysis confirmed comparable levels of DNA-PKcs in the heterozygous and kinase dead cell lines and a complete absence of protein in the knockout (Figure 5A-5B). We found that both the translocation frequency and the TL rate were dramatically increased in *DNA-PKcs^KD/-^*, but not *DNA-PKcs^-/-^* cells, compared to the parental *DNA-PKcs^+/-^*control (Figure 5C-5D). Further junction structure analysis of Cas9 and interchromosomal translocations indicated increased MMEJ only in *DNA-PKcs^KD/-^* and not in *DNA-PKcs^-/-^* cells (Figures 5E, S9C-S9D). Collectively, we conclude genetic DNA-PKcs kinase inhibition, similar to robust chemical DNA-PKcs inhibition, enhances translocations that are repaired more frequently by MMEJ.

**Figure 5.**
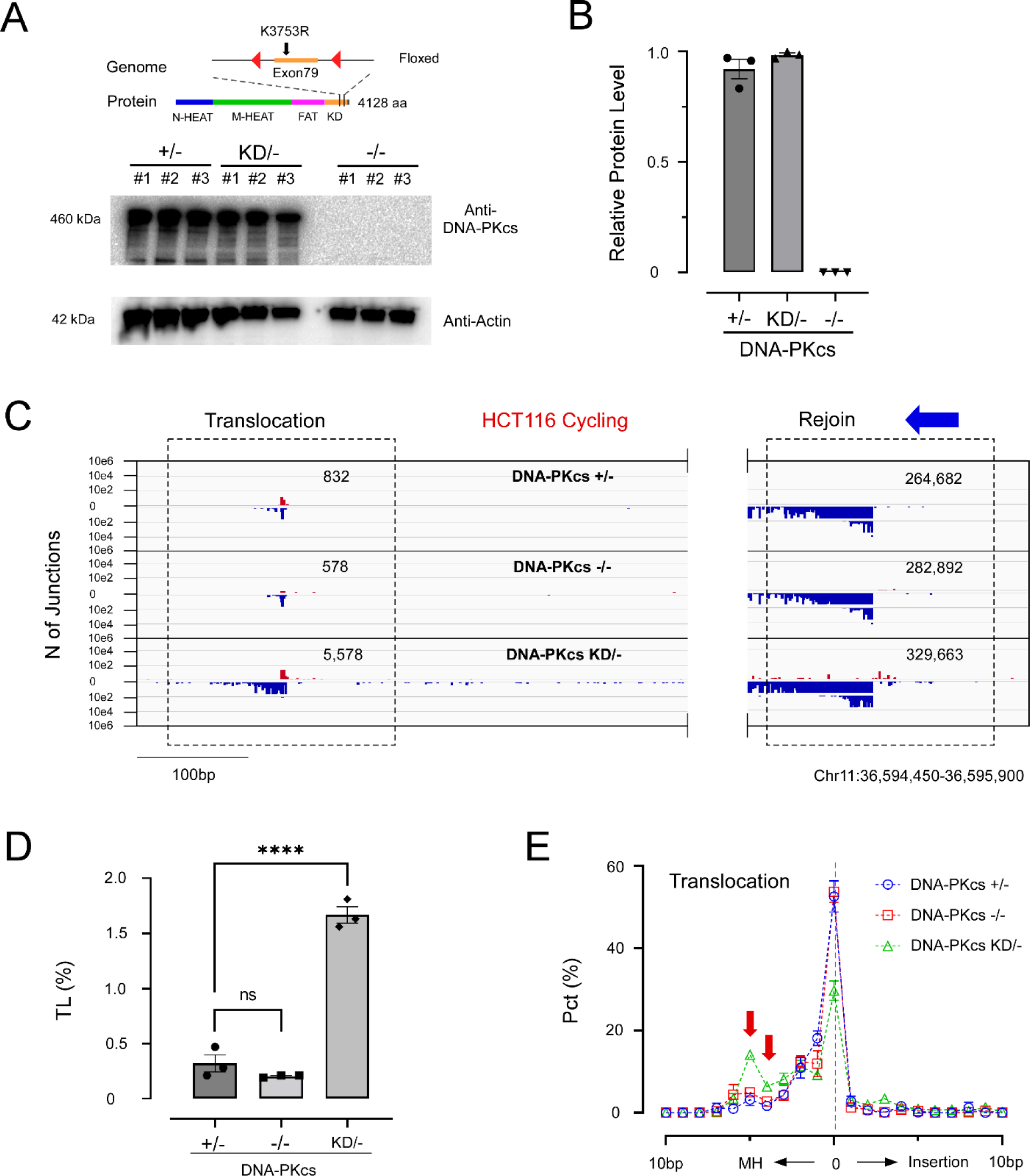
MH-mediated chromosome translocation increased in DNA-PKcs kinase dead HCT116 cells. (**A-B**) The exon79 of DNA-PKcs was deleted by flox-cre system and the kinase dead was generated by point mutation on K3753R. The western blot showed comparable level of DNA-PKcs protein in *DNA-PKcs ^+/-^* (control) and *DNA-PKcs ^KD/-^* but absent in *DNA-PKcs ^-/-^* (A) and the quantification was provided with error bar (B). (**C**) IGV translocation (RAG1D; left) and rejoin (RAG1L; right) plots of HCT116 cycling cells treated with genotypes including *DNA-PKcs ^+/-^* (control), *DNA-PKcs ^-/-^*and *DNA-PKcs ^KD/-^*. RAG1L bait priming (blue arrow) is indicated. (**D**) TL rates for *DNA-PKcs ^+/-^*, *DNA-PKcs ^-/-^* and *DNA-PKcs ^KD/-^* conditions were evaluated one-way ANOVA with ****p< 0.0001 and ns (no significance). (**E**) Translocation junction structure distributions of the above experiments with red arrows highlighting the increased MH utilization in *DNA-PKcs ^KD/-^* HCT116 cells.

### End Joining Pathway Choice on V(D)J Recombination is Determined by DNA-PKcs Protein and Kinase Status

With the above observations in mind for human cells, we wanted to discern NHEJ/A-EJ outcomes from DNA-PKcs inhibition versus deficiency in mouse *v-Abl* pro-B cell lines which undergo G1/G0 arrest and V(D)J recombination of hairpin sealed coding ends, and separately for blunt signal ends. We focused analysis on the Igκ light chain locus Vκ-Jκ pairing and cleavage of >100 functional Vκ-gene segments with four Jκ-gene segments by the RAG endonuclease; we used HTGTS-V(D)J-seq employing all functional Jκ coding end baits (Jκ1CE, Jκ2CE, Jκ4CE, and Jκ5CE) to assess DNA-PKcs perturbations (DPKi #2 (20μM) or DNA-PKcs exon 1 knockouts – *DNA-PKcs^-/-^* #1 and *DNA-PKcs^-/-^* #2) (Figure S10A) relative to wild-type (*WT*; NHEJ) and *Ku70^-/-^* (A-EJ) backgrounds^29^. For reference, the Jκ1 coding end bait in WT cells robustly forms coding joins (Figure 6A).

**Figure 6.**
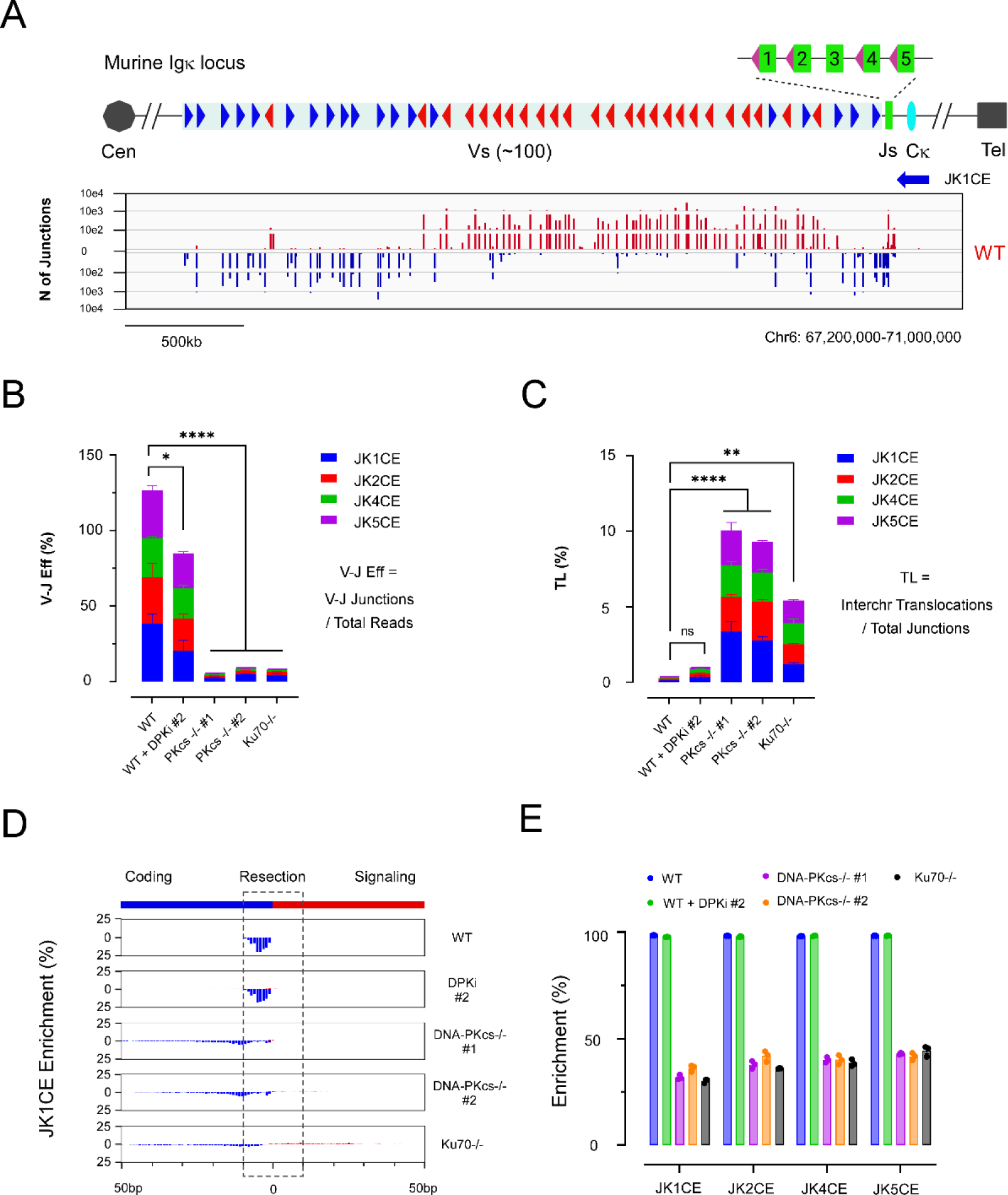
Jκ Coding Ends are Repaired by a Translocation-based A-EJ Mechanism Without DNA-PKcs. (**A**) Murine Igκ locus and representative junction profile of *WT v-Abl* cell Vκ-Jκ coding end recombination from the Jκ1 coding end (Jκ1CE) HTGTS bait. Resulting deletions (blue bars) and inversions (red bars) are directed by the RSS orientation. (**B**) V-J Eff (number of Vκ-Jκ junctions divided by the total reads in each library) is indicated for all four functional JκCE baits (Jκ1CE in blue, Jκ2CE in red, Jκ4CE in green, and Jκ5CE in magenta) in stacked bars. Combined values of *WT* (126.57 ± 17.81%), *WT* +DPKi #2 (84.82 ± 12.53%), *DNA-PKcs^-/-^* #1 (5.00 ± 0.34%), *DNA-PKcs^-/-^* #2 (8.65 ± 0.25%), and *Ku70^-/-^*(7.86 ± 1.74%) were evaluated by one-way ANOVA plus Dunnett’s posttest. (**C**) Interchromosomal (interchr) translocation fractions (TL Rate; junctions outside chr6 divided by the total junctions) for all four baits are shown in stacked bars. Combined values of *WT* (0.29 ± 0.04%), *WT* +DPKi #2 (0.95 ± 0.13%), *DNA-PKcs^-/-^* #1 (10.05 ± 1.33%), *DNA-*PKcs^-/-^ #2 (9.31 ± 0.63%), and *Ku70^-/-^* (5.41 ± 0.37%) were evaluated by one-way ANOVA plus Dunnett’s posttest. (**B,C**) *p<0.05; **p<0.01; ****p<0.0001; ns, not significant (**D**) Resected junction plots of the pooled Vκ gene fragment break sites (Jκ1CE bait) within a 100 bp window. RAG breakpoints are at position 0, flanked by coding (blue) and signal (red) sequences, with a 10bp resection cutoff (dashed box) indicated. (**E**) Junction enrichment within 10 bp of the DSB is quantified and shown for all baits. (**B-E**) Experiments were independently repeated three times with SEM.

We measured the Vκ-Jκ recombination efficiency (V-J Eff) and interchromosomal translocation fraction (TL) by normalizing Vκ region junctions against total reads and interchromosomal junctions against the total junctions recovered, respectively (Table S4). The recombination efficiency was decreased by >10-fold for *DNA-PKcs^-/-^* #1/2 and *Ku70^-/-^* cells compared to the *WT* control; *WT* +DPKi #2 cells were moderately depressed (Figure 6B), owing to the incomplete kinase inhibition. A corresponding reverse trend was found for the translocation fraction where DNA-PKcs deletion and inhibition increased the interchromosomal translocation rate (Figure 6C), suggestive of the utilization of A-EJ in the absence of DNA-PKcs. Junctions enriched around the ± 10bp coding/signal RAG incision sites of functional Vκ gene segment regions were quantified; unlike *WT* and *WT* + DPKi, which were not resected beyond 10bp, DNA-PKcs deletion reduced the enrichment by ∼70%, similar to *Ku70^-/-^* (Figures 6D-6E). Junction structures of Vκ-Jκ recombinants from JκCE baits revealed some distinguishing patterns. First, although common to all comparisons were the notable peak of direct joints, *WT* and *WT* +DPKi cells displayed the most (∼20%) insertions (<4 bp), consistent with NHEJ utilization; this was followed by *DNA-PKcs^-/^*^-^ #1/2 clones with an intermediate insertion fraction, while *Ku70^-/-^* possessed very few insertions. Second, while *WT* and *WT* +DPKi cells displayed fewer short MHs (1-3bps) than *Ku70^-/-^* cells, *DNA-PKcs^-/-^*#1/2 clones consistently tracked along the same *Ku70^-/-^* MH enrichment path for all baits (Figures S10B-S10E). Collectively, the coding end data suggest the absence, but not the inhibition, of DNA-PKcs has dramatic effects on V(D)J recombination and chromosome translocation, and the residual joining in the absence of DNA-PKcs harbors partially overlapping features of NHEJ and A-EJ.

Although signal-signal joining does not require DNA-PKcs,^62^ the consequence of promoting A-EJ from JκCE baits suggests a reciprocal possibility for signal end baits. Thus, we utilized corresponding Jκ signal ends (Jκ1SE, Jκ2SE, Jκ4SE, and Jκ5SE) using the above analysis parameters for evaluation (Figures S11A-S11G). Similar to CE baits, *WT* +DPKi #2 had a moderate decrease in recombination efficiency, whereas efficiency was reduced by ∼5-fold in the *DNA-PKcs^-/-^* #1/2 and *Ku70^-/-^* clones (Figure S11A)(Table S5). The translocation rate was significantly increased in both DNA-PKcs inhibited and deficient cells but were not as severe as *Ku70^-/-^* cells (Figure S11B), supporting the notion of varied NHEJ utilization contributing to signal end baits.^63,64^ In this regard, junctions from DNA-PKcs inhibited or deficient backgrounds were still predominantly biased for signal end preys and were not substantially resected (Figure S11C), nor were the nearly exclusive direct dominant signal end joints affected (Figures S11D-S11F). It is noteworthy that Vκ junctions from the Jκ5SE bait for all backgrounds assayed appeared to be involved in secondary recombination events from fused RSSs^33,65^ since they displayed a striking loss in signal end bias, substantial increase in resection, and a junction structure pattern resembling coding end joining (Figures S11C, S11G).

Considering the partial inhibition of DPKi #2, we evaluated three DNA-PKcs catalytically dead *v-Abl* cell lines, *DNA-PKcs^KD/KD^* #C1, #C3 and #C3-1 (D3922A)^39^ and examined the genetic inhibition on V(D)J recombination (Figure S12A-S12F) using Jκ1CE and Jκ1SE baits. In general, *DNA-PKcs^KD/KD^* clones harbored even more functional overlap with *Ku70^-/-^* than *DNA-PKcs^-/-^* cells. Specifically, kinase dead clones were ∼1.5% Vκ-Jκ efficient for both bait ends, which was below *Ku70^-/-^* (∼3-7%) but above *Lig4^-/-^* (∼0.2-0.4%) (Tables S4, S5) efficiency. Interestingly, and consistent with kinase dead activity in human cells, *DNA-PKcs^KD/KD^* clones displayed the highest rate of interchromosomal translocation among the *v-Abl* cells assayed (∼5% versus ∼1.5% for *Ku70^-/-^* Jκ1 baits) (Figures S12A-S12B). Furthermore, the joining fidelity, which was still biased for signal ends without DNA-PKcs (Figure 8A), was also lost with a bias rate for both coding and signal ends similar to *Ku70^-/-^* (∼2 fold) (Figure S12C). However, V region resected joints displayed an intermediate level of enrichment relative to *WT* and *Ku70^-/-^*, with 70% and 80% junctions within ±10 bp window for Jκ1CE and Jκ1SE baits, respectively (Figure S12D); these resected joints were accompanied by a nearly 2-fold drop in direct repair relative to *WT* with increased MMEJ for both baits in a manner mostly resembling *Ku70^-/-^*(Figure S12E-S12F). We conclude DNA-PKcs is essential for maintaining the fidelity of Vκ-Jκ recombination by suppressing chromosome translocation.

### DNA-PKcs Promotes MMEJ in the Absence of *Lig4*

Using deeper sequencing and closer nested primer to bait in the *v-Abl* studies described here, we can recover 10x more junctions from *Lig4^-/-^* RAG DSBs than previously reported,^29^ yielding a total junction average of ∼1,800 and ∼1,400 per replication for JκCE and JκSE baits, respectively (Figure 7A; Tables S4 and S5). In contrast, *Lig4^-/-^Ku70^-/-^* total junctions averaged ∼10,000 and ∼12,000 per replication for the above baits (Tables S4 and S5), yielding >5-fold increase in Vκ-Jκ efficiency (Figures 7B and S13A) and were indistinguishable to *Ku70^-/-^ v-Abl* cells.^30^ Thus, we investigated whether DNA-PKcs blocks A-EJ in the *Lig4^-/-^* context like Ku70. Neither DNA-PKcs inhibition or deletion were found to increase Vκ-Jκ efficiency; in comparison, *Lig4^-/-^Ku70^-/-^*increased Vκ-Jκ efficiency by ∼10-fold (Figures 7B and S13A). DNA-PKcs inhibition and deficiency did not increase the already high translocation rate in *Lig4^-/-^* cells (relative to *WT*), as it does for *Lig4^-/-^Ku70^-/-^* coding end baits (Figures 7C and S13B). Overall, the data suggest DNA-PKcs does not block the Ku-independent A-EJ pathway to repair RAG DSBs.

**Figure 7.**
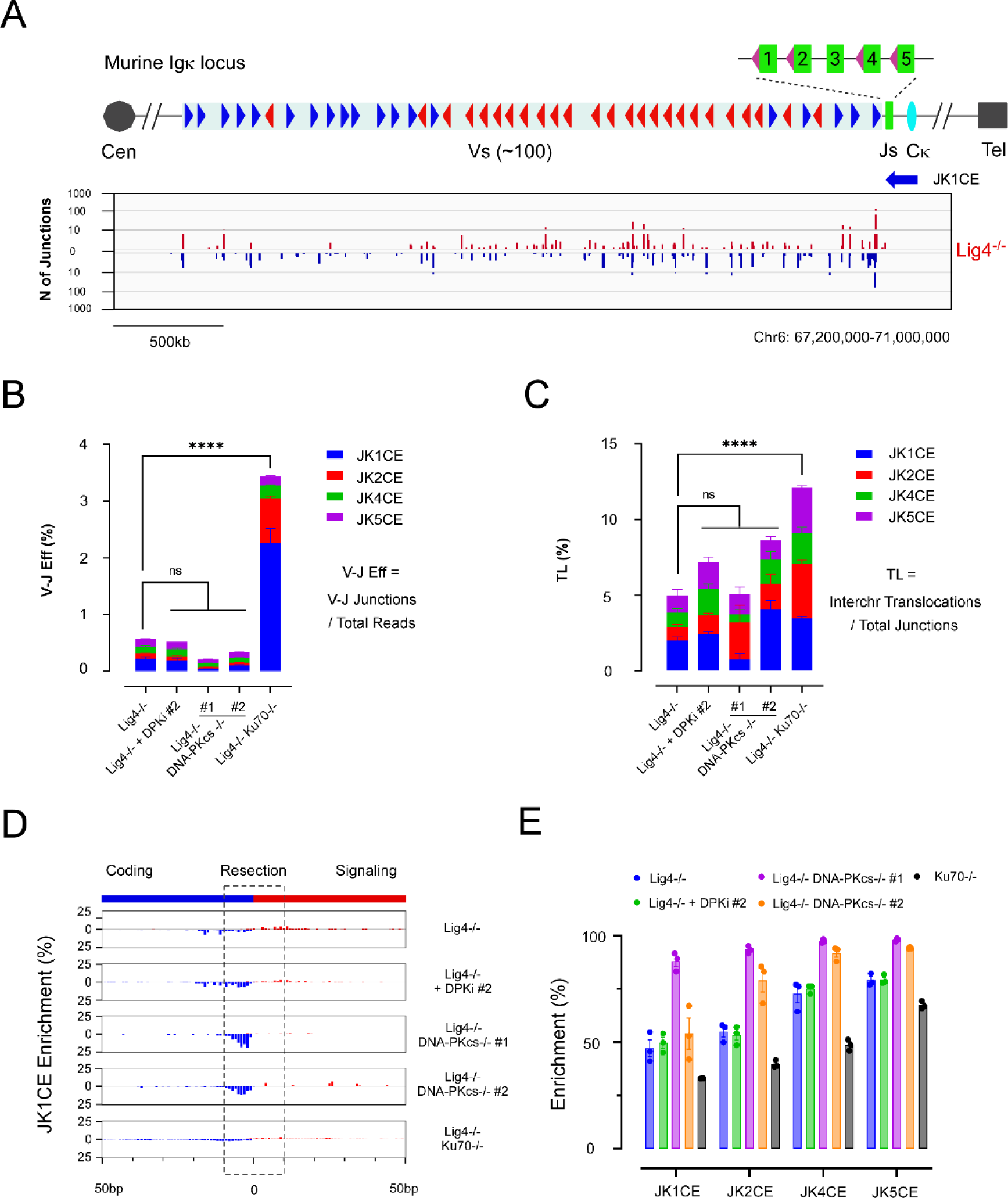
DNA-PKcs Promotes Resected A-EJ of Jκ Coding Ends Without DNA Ligase 4. (**A**) Representative profile of Vκ-Jκ coding ends recombination in *Lig4^-/-^ v-Abl* cells, using the Jκ1CE HTGTS bait. See Figure 5 legend for more details. (**B**) V-J Eff from the four functional Jκ coding end baits are indicated. Combined values of *Lig4^-/-^* (0.57 ± 0.05%), *Lig4^-/-^*+DPKi #2 (0.52 ± 0.07%), *Lig4^-/-^ DNA-PKcs^-/-^* #1 (0.21 ± 0.02%), *Lig4^-/-^DNA-PKcs^-/-^* #2 (0.33 ± 0.01%), and *Lig4^-/-^Ku70^-/-^*(3.44 ± 0.29%) were statistically distinguished using one-way ANOVA plus Dunnett’s posttest. (**C**) Normalized Translocation, TL Rate, are indicated for all four baits. Combined values of *Lig4^-/-^* (4.97 ± 0.22%), *Lig4^-/-^* +DPKi #2 (7.19 ± 0.53%), *Lig4^-/-^ DNA-PKcs^-/-^*#1 (5.09 ± 1.57%), *Lig4^-/-^ DNA-PKcs-/-* #2 (8.63 ± 1.19%), and *Lig4^-/-^Ku70^-/-^* (12.08 ± 0.48%) were distinguished using one-way ANOVA plus Dunnett’s posttest. (**B,C**) ****p<0.0001; ns, not significant. (**D**) Resected Vκ region junction plots from Jκ1CE bait are shown; see Figure 5 legend for more details. (**E**) Resected junction summary in all indicated conditions for the four JκCE baits. (**B-E**) Experiments were independently repeated three times with SEM indicated.

Investigation of resection and junction structures proceeded with Vκ-Jκ junctions. In general, *Lig4^-/-^*prey resected joints from both JκCE and JκSE baits was greater than *WT* but was less than *Ku70^-/-^*; resected joints were similarly increased in *Lig4^-/-^Ku70^-/-^*cells to a comparable level to *Ku70^-/-^* alone. DPKi #2 had no added effect on *Lig4^-/-^* cells (Figures 7D-7E and S13C), likely due to incomplete inhibition; though we observed only a modest increase in resected joints relative to *WT* with *DNA-PKcs^KD/KD^* clones (Figure S12D) which we infer to be stabilized by the presence of Lig4. Notably for JκCE baits, we observed a decreased range of resected joints with *Lig4^-/-^DNA-PKcs^-/-^* #1/2 clones compared to *Lig4^-/-^* alone, which aligns with recent studies demonstrating that DNA-PKcs promotes DSB resection.^66,67^ In JκSE baits, *Lig4^-/-^DNA-PKcs^-/-^*clones displayed varied effects on resection (Figure S13C). DNA-PKcs and Ku70 deletion also decreased MMEJ (>2bp) and correspondingly increased direct repair for all baits compared *Lig4^-/-^*control (Figures S14A-S14D and S13D-S13G). Collectively, these findings support a G1/G0-phase NHEJ variant A-EJ pathway that is revealed by Lig4 deficiency and reliant on DNA-PKcs for resection to promote MMEJ.

## Discussion

Elucidating the crosstalk between NHEJ and A-EJ mechanisms has been the subject of intense study in recent years. Our report adds further distinction between DNA-PKcs kinase versus structure functions as it relates to chromosome translocations and MMEJ. Specifically, we found the kinase activity of DNA-PK to be an important step in suppressing translocations, where translocations in human and mouse cells are normally distinguished by direct versus MH joints, respectively repaired by NHEJ and MMEJ.^22^ On one hand, our data show complete kinase inhibition led to a block in NHEJ and a rise in alternative repair mechanisms that leverage MMEJ to varying degrees. The microhomology increase with complete kinase inhibition in our human and mouse experiments is aligned with recent findings demonstrating increased MMEJ-mediated translocations^68^ or homology-directed repair^57^ upon robust DNA-PKcs kinase inhibition. On the other hand, we demonstrate incomplete chemical inhibition of DNA-PKcs leads to a shift in the balance between DSB rejoining and translocation and a drop in Vκ-Jκ joining fidelity (Figure 8A), therefore, enabling transient kinase inhibition as the driving mechanism to swap synapsed DNA end partners and re-engage NHEJ for efficient repair. In this regard, proximal and frequent DSBs would be highly selected end partners for translocation^30^ and, therefore, ideal for enhancing gene edited deletions. Thus, in addition to the 20 to 50-fold increased expression of DNA-PKcs in humans versus mice,^69,70^ kinase activity, likely related to autophosphorylation,^39,63^ is another contributing factor in the differential manifestation of chromosome translocations between humans and mice. While yet to be tested in mice, a recent report of a novel ATM-mediated phosphorylation site at the extreme C terminal end of DNA-PKcs (T4102) which helps to enhance overall DNA-PKcs functions, specifically stabilizing the DNA-PKcs-Ku interaction and recruitment of processing factors,^71^ may bring further insight into NHEJ transition and translocation mechanisms. Despite the end-joining choice, DNA-PKcs suppresses chromosome translocation in both mammals.

**Figure 8.**
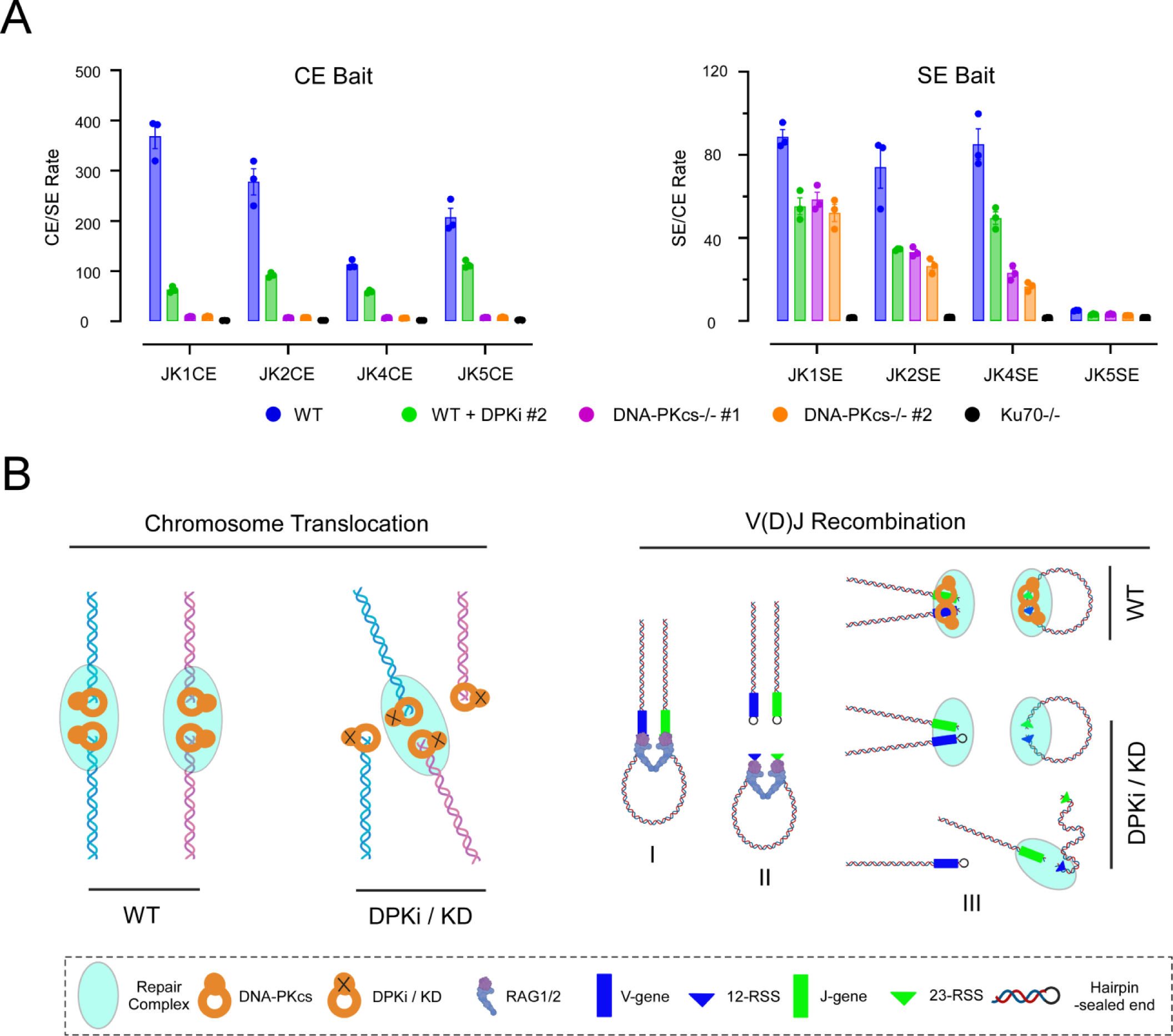
DNA-PKcs Suppresses Illegitimate Chromosome Rearrangement. (**A**) Coding end and signal end bias, relative to the bait used, are progressively lost with DNA-PKcs perturbations. Note *DNA-PKcs^-/-^* is more severely affected than DPKi treatments. (**B**) Model illustrating how DNA-PKcs prevents deleterious chromosome rearrangement.

Although the classic NHEJ transition mechanism involves MRE11 endonuclease/exonuclease activity and eviction of Ku70/80 or with DNA-PKcs,^67^ this mechanism is typically viewed through the lens of homologous recombination. Our data support the recent growing evidence of a transition mechanism into using MMEJ that involves DNA-PKcs and Lig4 which could provide insight into end joining pathway choice. Specifically, we found that DNA-PKcs possesses opposing DNA end resection functions for G1/G0-phase RAG DSBs with the presence of Lig4 suppressing, and the absence of Lig4 promoting, resection. Though suppressing resection aligns with excluding A-EJ, promoting resection was surprising and through a mechanism that is not fully understood.^66^ It is possible that the immature DNA-PK synapsis at DNA ends^64^ provides a favorable platform for recruiting DNA polymerases, nucleases, and other factors to support an Artemis & resection-dependent NHEJ mechanism that was previously described in G1, but not G2, phase.^72,73^ In this regard, ATM kinase can activate Artemis in the presence of an inhibited DNA-PKcs^39,74^ and could be used to re-engage NHEJ-mediated ligation in the presence of Lig4. Thus, our data suggest the signal to stop end process cycling involves *Lig4*, likely with other NHEJ short range complex components.^8,9,63,64^ Whether this persistent DNA-PKcs-dependent resection in the absence of Lig4 is related to the Artemis dependent mechanism, and whether this process more broadly represents an NHEJ-variant mechanism to explain MMEJ-mediated translocations in cycling cells, remain to be determined.

Collectively, our findings support a model (Figure 8B) that DNA-PKcs suppresses chromosome translocations in both human and mouse cells, albeit by different repair path mechanisms, where “adrift” unpaired DNA ends, due to kinase inhibition, enable greater opportunities to form translocations if the kinase inhibition efficacy is low (left panel). Additionally, DNA-PKcs safeguards the integrity of V(D)J recombination (right panel), by playing an essential role after Vκ and Jκ RSSs undergo pairing (I) and cleavage (II) by RAG. In the postcleavage complex, DNA-PK is necessary for rapid coding-coding recombination through NHEJ and suppressing hybrid joints (coding-signal) through its kinase activity, perhaps in collaboration with ATM.^74,75^.

Conversely, without DNA-PKcs, hairpin-sealed ends are not resolved efficiently, leading to A-EJ-mediated coding-coding and hybrid joints, but leaving signal-signal end recombination by NHEJ mostly intact (III). The absence of Lig4 creates further opportunity for DNA-PKcs to promote resection and MMEJ, an activity that is more readily apparent on coding ends where hairpin opening creates an overhang that necessitates end processing.

Chromosome translocation is a driver for many cancers including leukemias and lymphomas. Therefore, it is crucial to balance the fitness gains of an efficient V(D)J recombination relative to the deleterious impact of translocations. Previous studies demonstrated that chromosome translocations in cycling murine cells are mediated by A-EJ,^20,21^ and A-EJ is often suppressed by NHEJ. ^22,29,50^ In agreement, we found DNA-PKcs kinase dead in both humans and mice synergizes translocations, requiring A-EJ mechanisms, which include MMEJ, to complete their repair. We conclude DNA-PKcs is essential for genome stability and maintaining the fidelity of V(D)J recombination by suppressing chromosome translocation and regulating end joining pathway commitment.

## Data Availability

HTGTS data of human and mouse data were submitted to Gene Expression Omnibus (GEO) and publicly available with the following accession numbers: Human GSE232940; Mouse GSE232944. Other data for determining the boundary of topology-associated domains are publicly available in GEO: Chip-seq for CTCF in K562 cells (GSM749733, GSM935407 and GSM733719); Chip-seq for Rad21 in K562 cells (GSM935319); Chip-seq for CTCF in HEK293 cells (GSE91917 and GSM749687).

## Acknowledgements

We thank the Bassik laboratory for the ACOC gRNA library, Ramesh Nair from Stanford Center for Genomics and Personalized Medicine (SCGPM) for his advice on data analysis, and the Stanford Cancer Institute core facilities. We also thank the Davis (UT Southwestern),^38^ Zha (Columbia),^39^ and Meek (U Michigan)^61^ laboratories for providing reagents.

