## Supplementary Figures for "DNA-PKcs Suppresses Illegitimate Chromosome Rearrangements"

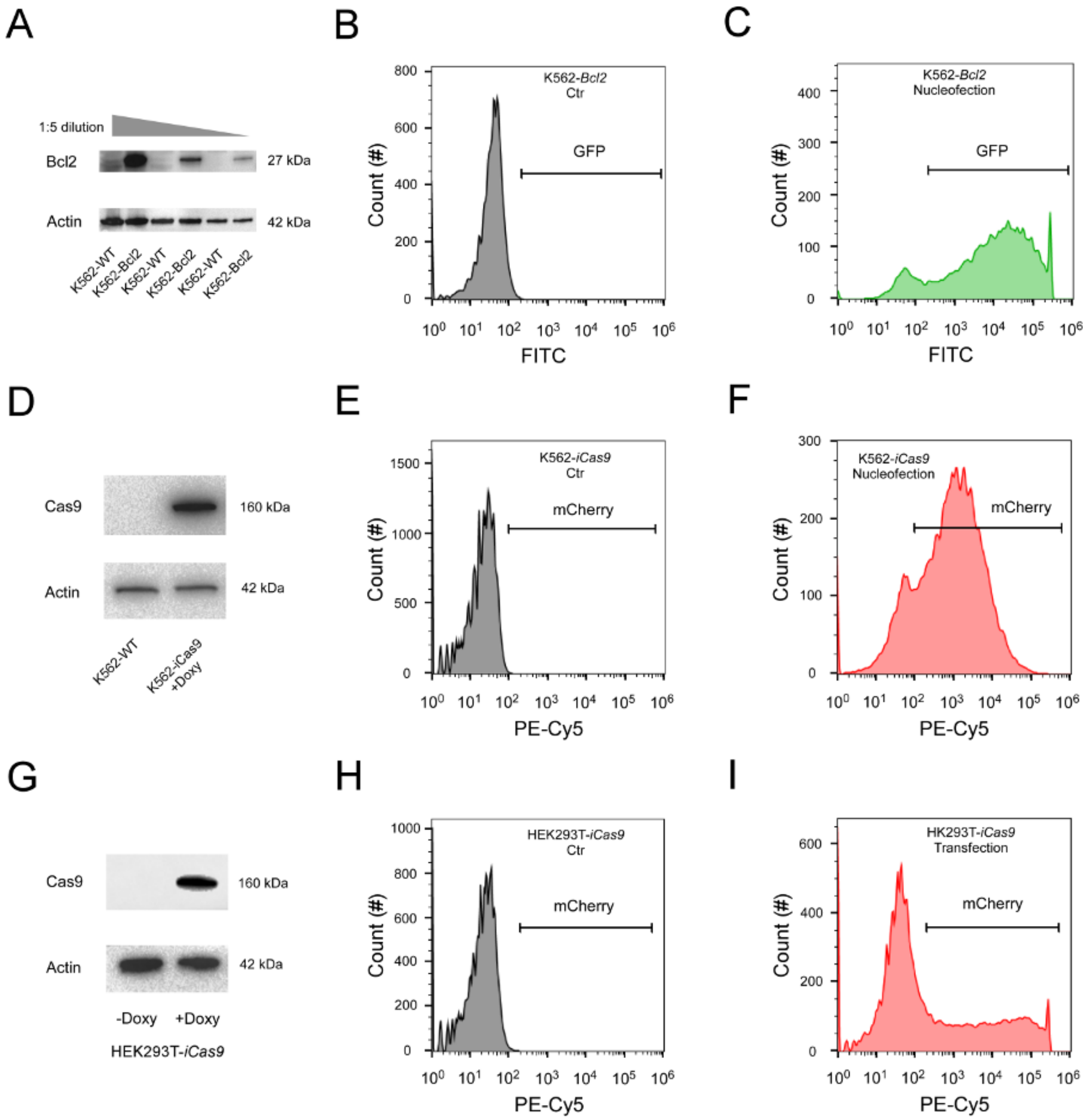


**Supplementary Figure S1. Verification of cell lines and quantification of transfection efficiency.** (**A**) Bcl2 expression levels were quantified by Western blot in *K562-Bcl2* and its parental cell *K562-WT* using a Bcl2 antibody. The input was set as 1:5 dilution gradients and controlled by the housekeeping protein Actin. (**B-C**) Transfection efficiency was evaluated by combining all nucleofection assays using pX330-Cas9-gRNA with a control vector pMax-GFP and quantified by flow cytometry. The representative histograms of nucleofection efficiency without nucleofection (**B**) and with pX330-Cas9-gRNA (RAG1D/L) plus pMax-GFP (**C**) are shown, with the horizontal bar indicating FITC. (**D**) The Cas9 expression level in *K562-iCas9* (with the insertion of the Cas9-Flag-T2A -GFP-NeoR cassette) upon doxycycline (+Doxy) induction was examined by Western blot using an anti-flag antibody. No Cas9 level was detected in its parental cell line *K562-WT*. (**E-F**) Representative figures of nucleofection efficiency in the conditions without nucleofection control (**E**) and in the condition with pMCB320-gRNA in *K562-iCas9* cells, as pMCB320-gRNA contains a mCherry reporter that can serve as an indicator of nucleofection efficiency. (**G-I**) Same as (**D-F**) but for *HEK293T-iCas9* cells by lipofectamine-mediated transfection (with insertion of Cas9-Flag-T2A -GFP-BlastR cassette).

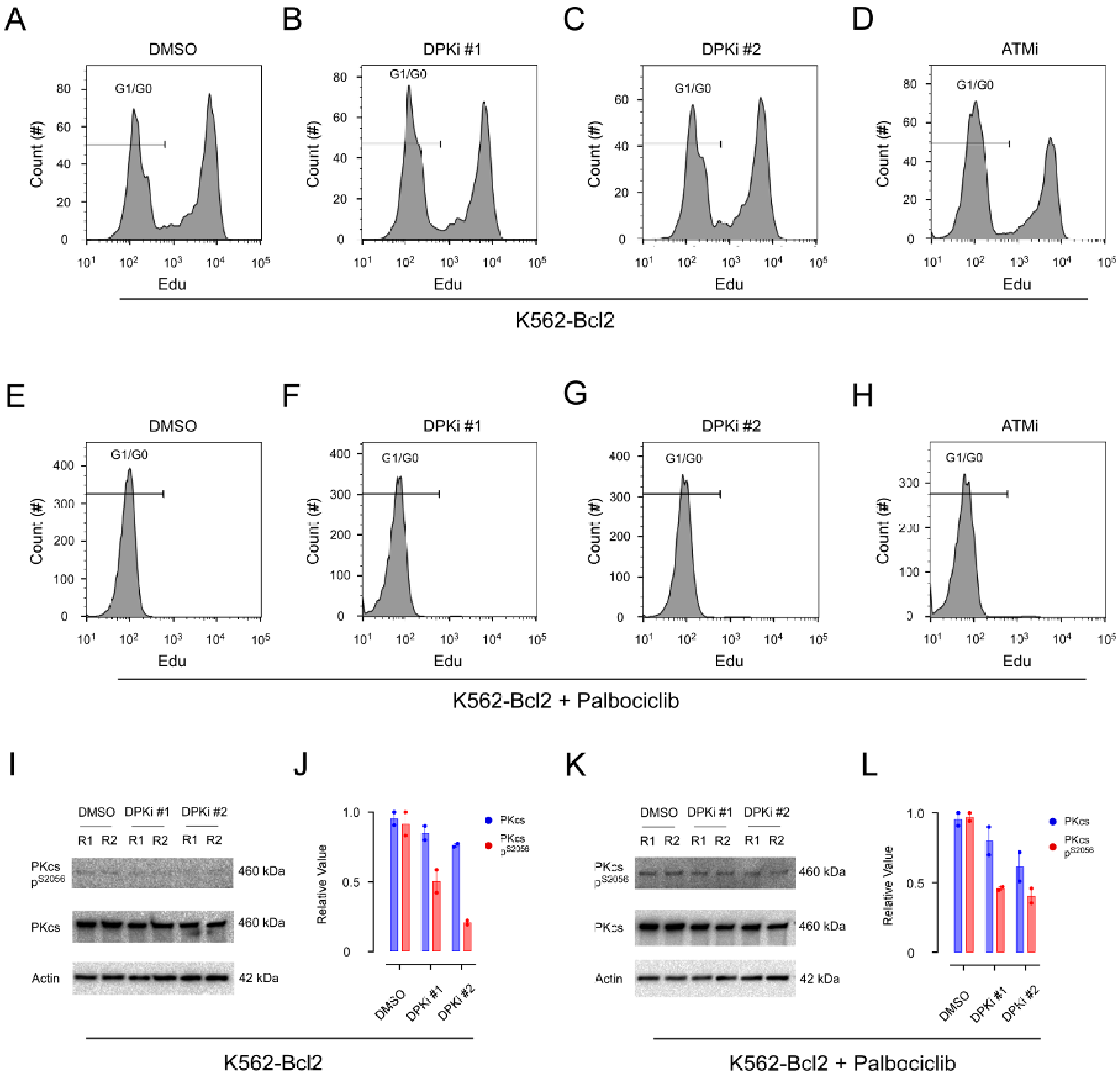


**Supplementary Figure S2. Cell cycle analysis following treatment with DNA-PKcs and ATM inhibitors.** (**A-D**) Flow cytometry analysis of *K562-Bcl2* cell cycle distribution after treatment with DMSO (**a**), DPKi #1 (**B**), DPKi #2 (**C**), and ATMi (**D**). Cells were incubated with Edu, clicked with FITC-488, and quantified by flow cytometry. The G1/G0 population is highlighted. (**E-H**) Same as (a-d) but for Palbociclib in combination with the inhibitors. Results indicate that the cell cycle was not affected by treatment with DNA-PKcs inhibitors. All experiments were independently repeated at least three times. (i-j) The protein levels of DNA-PKcs and its phosphorylation state at S2056 ($p^{S2056}$) were analyzed by western blotting two days after nucleofection with pX330-RAG1D/L in *K562-Bcl* cells. The cells were treated with DMSO, DPKi #1, or DPKi #2 (i), and the experiments were independently repeated twice (R1 and R2). The protein levels of DNA-PKcs and $p^{S2056}$ were normalized to Actin and presented in panel j. (k-l) Similarly to (i-j), the protein levels of DNA-PKcs and $p^{S2056}$ were examined after treatment with Palbociclib. The experimental procedures were conducted as described for (i-j).

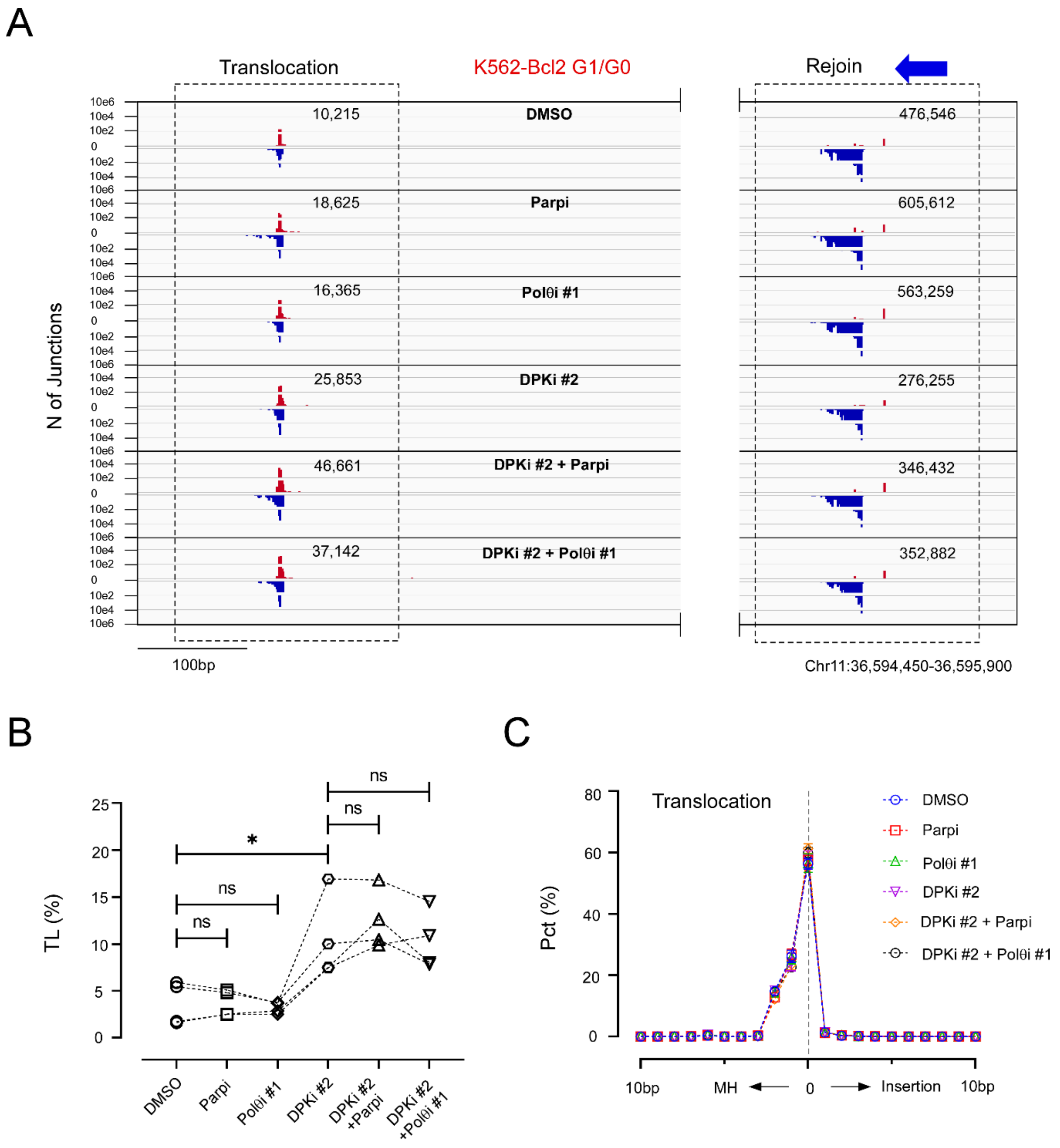


**Supplementary Figure S3. Effects of A-EJ pathway inhibitors, Olaparib and Novobiocin, on chromosome translocation.** (**A**) *K562-Bcl2* cells arrested by Palbociclib 50 mM and treated with DMSO, Parpi (Parp1 inhibitor, Olaparib, 10μM), Polθi #1 (Polθ inhibitor, Novobiocin, 100 μM) with/without DPKi #2, with the junctions of translocation and rejoin captured by HTGTS with RAG1L bait (blue arrow) and plotted by IGV. (**B**) The values of normalized translocation (TL) under conditions of DMSO, Parpi, and Polθi #1 with/without DPKi #2, which were evaluated by ratio paired T-test with *$p < 0.05$ and ns (no significance).. The dashed line links the same batch of measurements. (**C**) The repair patterns of translocations, including microhomology (MH), insertion, and direct repair, in a 20 bp window in percentage (Pct, %). All experiments were independently repeated four times, and the standard error of the mean (SEM) is provided.

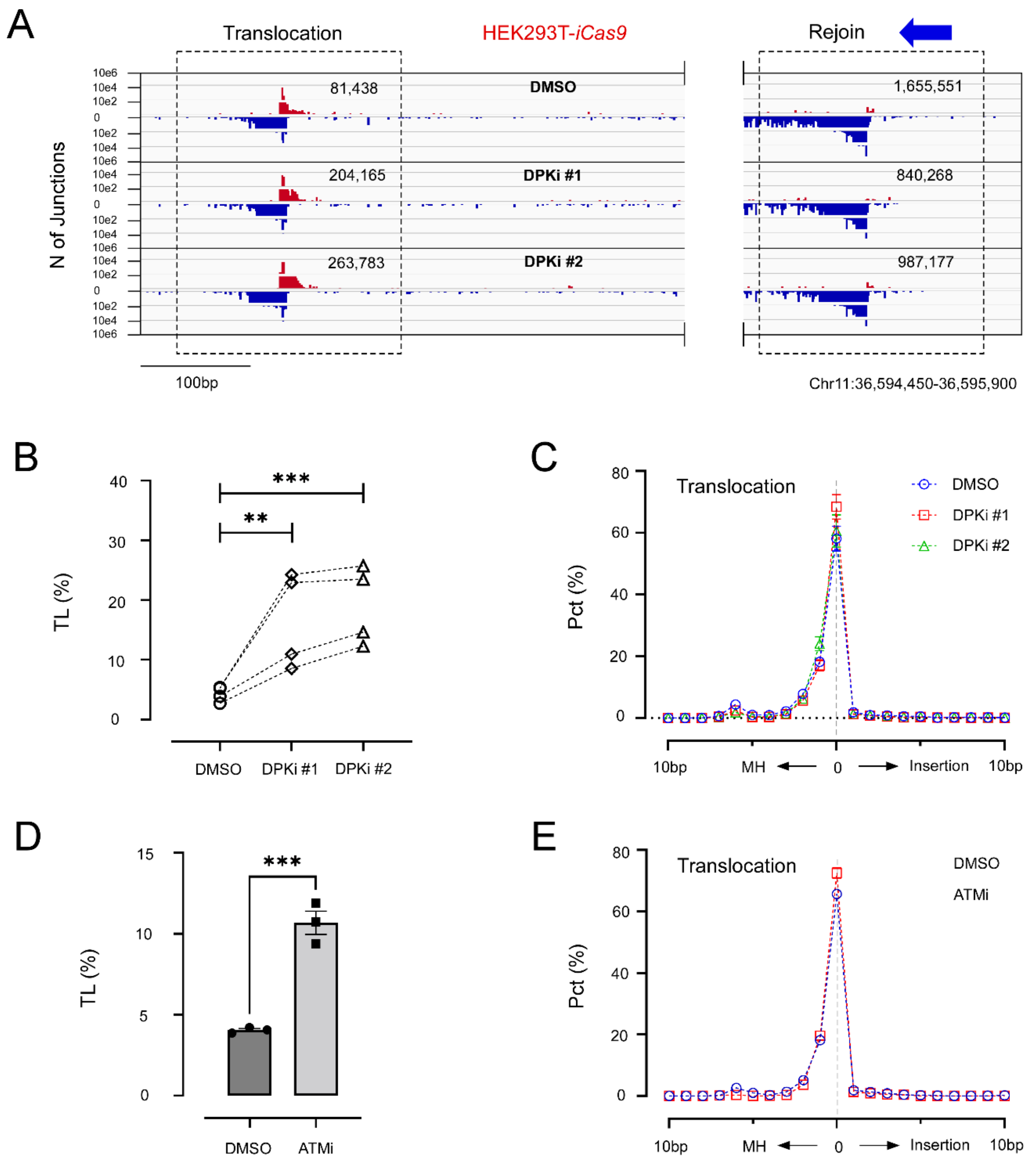


**Supplementary Figure S4. DNA-PKcs inhibition promotes NHEJ-mediated chromosome translocation in *HEK293T-iCas9* cells.** (**A**) Two DSBs were generated in *HEK293T-iCas9* cells by transfection of twinned pMCB320-gRNA (RAG1D + RAG1L). The resulting translocations and rejoin events in conditions with DMSO, DPKi #1 and DPKi #2 were captured by HTGTS and shown as an IGV plot. The blue arrow indicates the bait site. (**B**) The normalized translocation levels (TL) for DMSO, DPKi #1, and DPKi #2 were evaluated by a paired ratio T-test with **$p<0.01$ and ***$p<0.001$. The dashed lines linking the data in the same batch. (**C**) The repair patterns of translocations, including microhomology (MH), insertion, and direct repair, in a 20 bp window, presented as percentages (Pct, %). (**D**) Same as (B) but for ATMi and experiment performed in one batch and evaluated by standard T-test with ***$p<0.001$. (**E**) Same as (C) but for ATMi. All experiments were independently repeated at least three times (n=3), and the standard error of the mean (SEM) is provided.

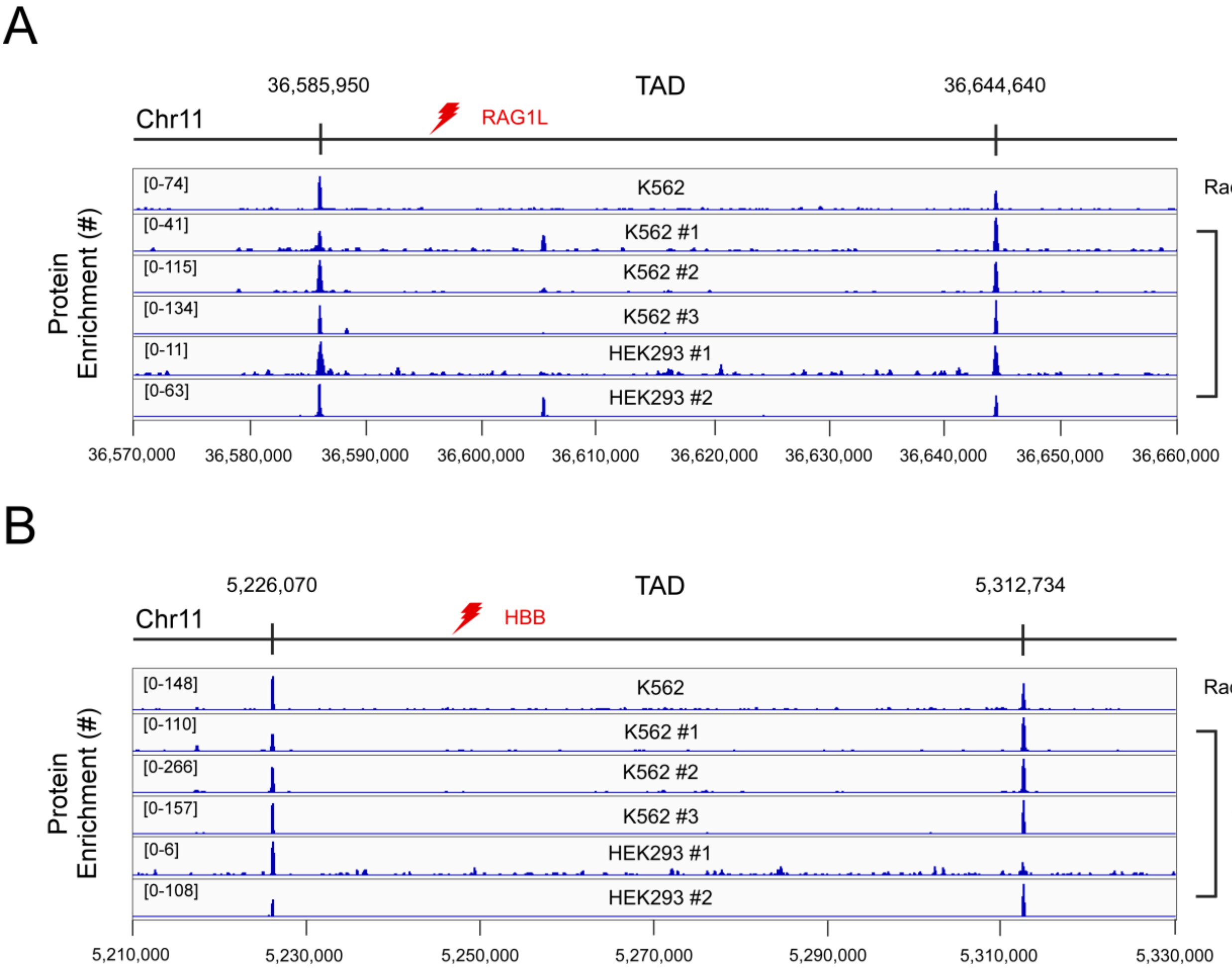


**Supplementary Figure S5. Determining the boundaries of the topologically associated domain (TAD) for RAG1L and HBB baits.** (**A**) The boundary of the TAD encompassing the RAG1L bait site (red lightning bolt) on chromosome 11 was determined by aligning Rad21 and CTCF Chip-Seq data obtained from K562 and HEK293 cells against the hg19 genome. The convergence of CTCF motifs, along with the enrichment of CTCF and cohesion Rad21 proteins, identified the sites at 36,585,950 and 36,644,640 as the boundary of the TAD encompassing RAG1L. (**B**) The TAD encompassing the HBB bait site (red lightning bolt) was determined using a similar approach, resulting in the identification of the region chr11:5,226,070-5,312,734 as the TAD boundary.

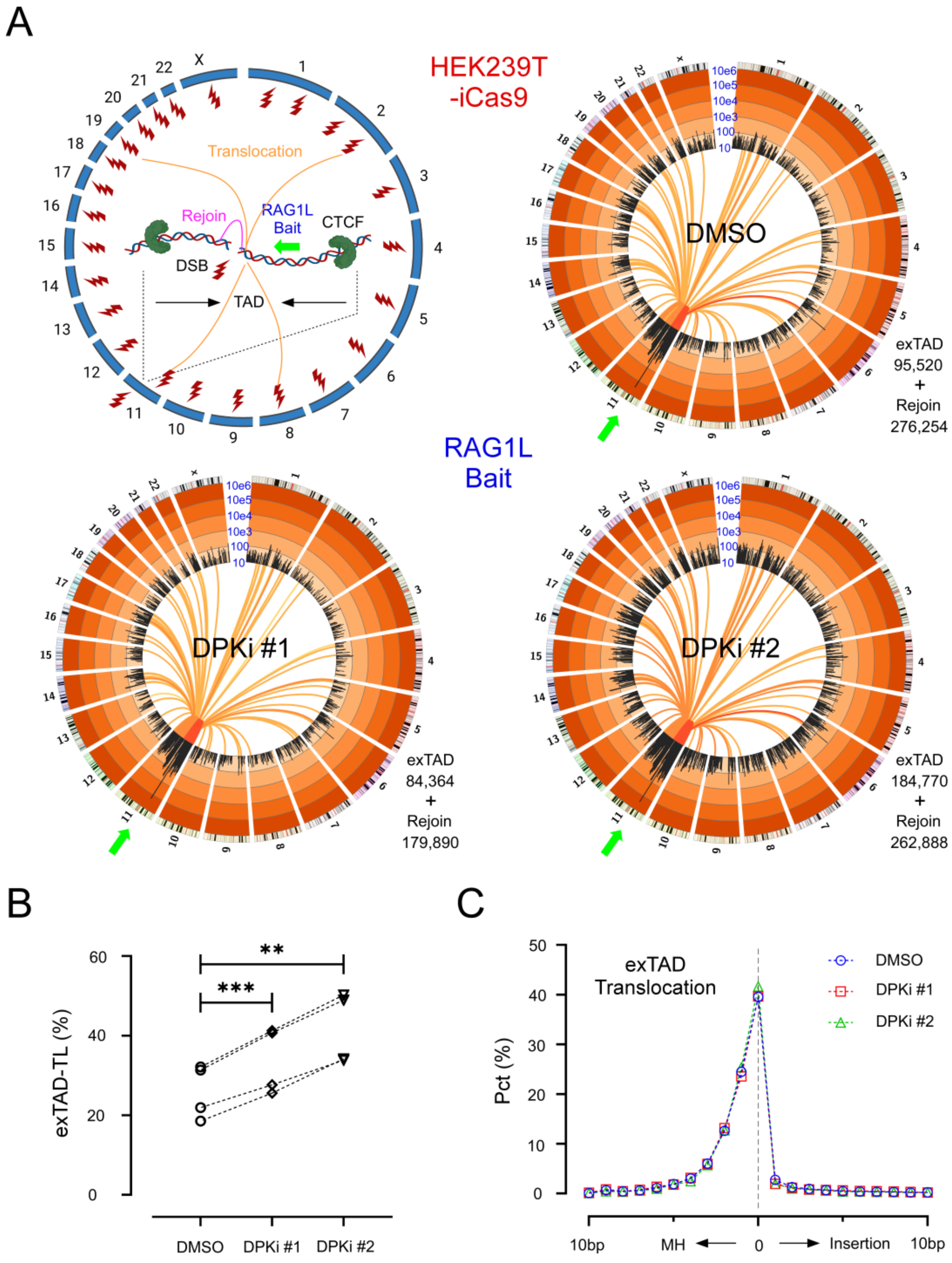


**Supplementary Figure S6. DNA-PKcs inhibition increases genome-wide chromosome translocation in *HEK293T-iCas9* cells.** (**A**) To generate double-strand breaks (DSBs) in genome-wide manner in *HEK293T-iCas9* cells, ACOC library containing ~30,000 gRNA guides (represented as red lightning bolts) was employed. The RAG1L gRNA was used as bait for HTGTS, shown as a green arrow. The translocations and rejoins are depicted by orange and magenta lines, respectively. The bait site is flanked by CTCF binding sites, forming a Topology-Associated Domain (TAD, Chr11: 36,585,950-36,644,640). The genome-wide chromosome translocation profiles for DMSO, DPKi #1, and DPKi #2 conditions are shown as circos plots. (**B**) The number of translocations outside the TAD region (exTAD-T) is normalized to the number of rejoins as exTAD-TL. The exTAD-TL changes between DPKi #1, DPKi #2 and DMSO were evaluated by a ratio paired T-test with $^{**}p < 0.01$ and $^{***}p < 0.001$. Dashed lines link the data obtained from the same batches. (**C**) The repair pattern of exTAD-T is presented in a 20 bp window as a percentage (Pct, %). All experiments were independently repeated four times, and the standard error of the mean (SEM) is provided.

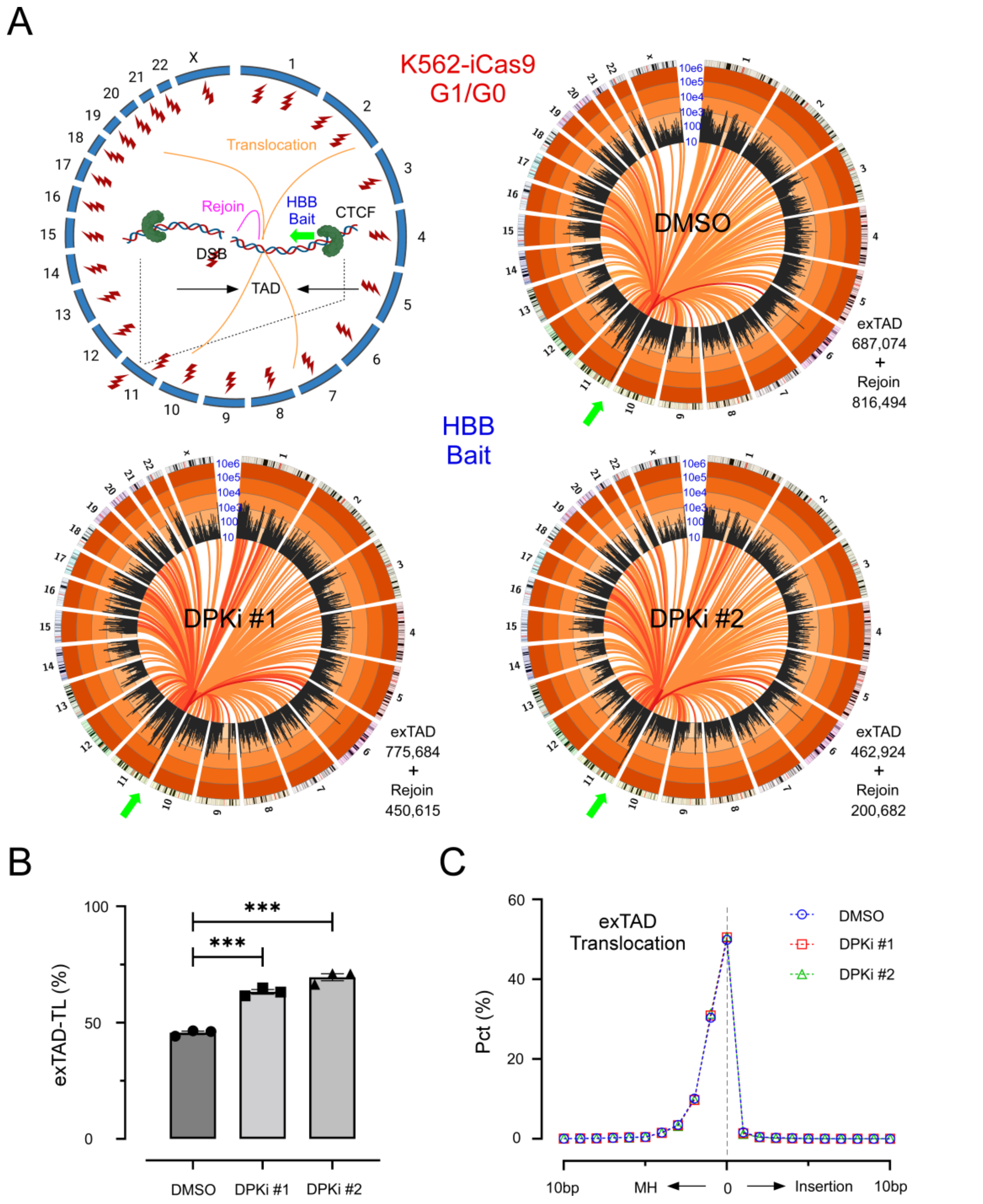


**Supplementary Figure S7. DNA-PKcs inhibition increases genome-wide chromosome translocation in *K562-iCas9* cells using HBB bait.** (**A**) To generate double-strand breaks (DSBs) in genome-wide manner, we used ~30,000 ACOC gRNA guides (represented as red lightning bolts) in G1/G0-arrested *K562-iCas9* cells. The HBB gRNA was used as a bait for high-throughput genome-wide translocation sequencing (HTGTS, shown as a green arrow). The translocations and rejoins are depicted by orange and magenta lines, respectively. The bait site is flanked by CTCF binding sites, forming a Topology-Associated Domain (TAD, Chr11: 5,226,070-5,312,734). The genome-wide chromosome translocation profiles for DMSO, DPKi #1, and DPKi #2 conditions are shown as circos plots. (**B**) The number of translocations outside the TAD region (exTAD-T) is normalized to the number of rejoins as exTAD-TL. The exTAD-TL changes between DMSO (45.67 ± 0.72 %, n=3), DPKi #1 (63.25 ± 0.98 %, n=3), and DPKi #2 (69.52 ± 1.51 %, n=3) were evaluated by unpaired T-test with ***$p < 0.001$. (**C**) The repair pattern of exTAD-T is presented in a 20 bp window as a percentage (Pct, %). All experiments were independently repeated three times, and the standard error of the mean (SEM) is provided.

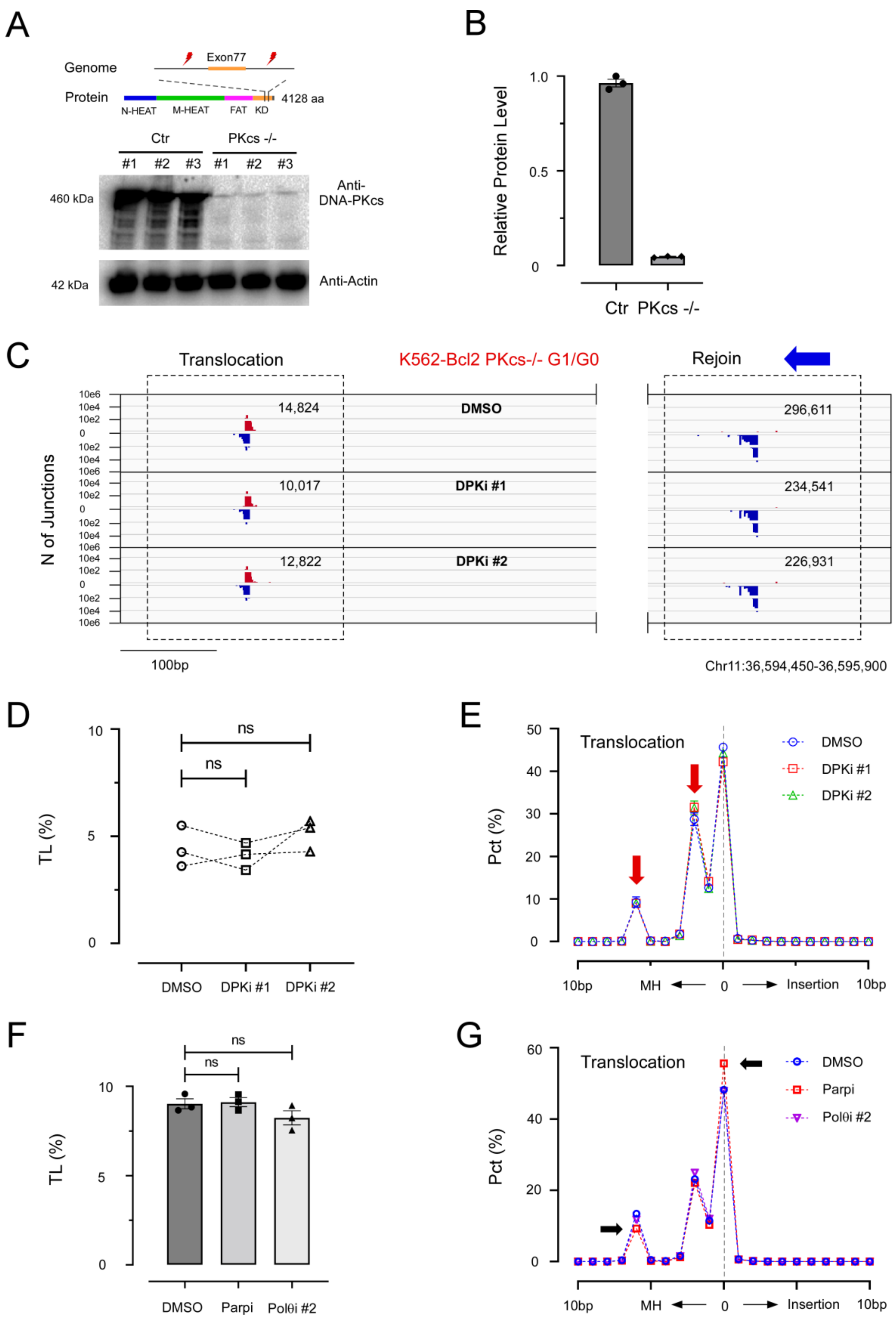


**Supplementary Figure S8. Deletion of the DNA-PKcs kinase domain increases MH-mediated translocation.** (**A**) A schematic representation of the experimental setup using twined Cas9:gRNAs (red lightning bolts) to delete exon 77 of

DNA-PKcs (yellow) in the *K562-Bcl2* genome. This deletion resulted in the loss of functional DNA-PKcs kinase domain. Western blot analysis using a DNA-PKcs antibody revealed reduced expression of truncated DNA-PKcs in *K562-Bcl2 DNA-PKcs*$^{-/-}$ (*PKcs*$^{-/-}$) cells compared to the parental cell line, *K562-Bcl2* (Ctr). Each condition contained three independent western blot samples, #1, #2 and #3. (**B**) The protein level of DNA-PKcs and its kinase domain truncated version in (A) was quantified by setting the lane of DNA-PKcs in Ctr #1 as 1, and normalize the rest, respectively. (**C**) IGV translocation (RAG1D; left) and rejoin (RAG1L; right) plots of G1/G0-arrested *K562-Bcl2 PKcs*$^{-/-}$ cells treated with DMSO, DPKi #1, and DPKi #2. RAG1L bait priming (blue arrow) is indicated. (**D**) TL rates for DMSO, DPKi #1, and DPKi #2 conditions were evaluated by ratio paired T-test, generating geometric mean of ratios and SEM of log for DPKi#1 vs DMSO (0.924 ± 0.049) and DPKi#2 vs DMSO (1.159 ± 0.039). Dashed lines link the data obtained from the same batch of experiments. (**E**) Translocation junction structure distributions of the above experiments with red arrows highlighting the increased MH utilization. (**F**) TL rates for DMSO, Parpi and Polθi #2 on top of G1/G0-arrested *K562-Bcl2 PKcs-/-* cells. The differences were evaluated by one-way ANOVA. (**G**) Same as (F) but for Parpi and Polθi #2 with black arrow highlighting the changes upon Parpi treatment. All experiments were independently repeated three times, and SEM are indicated.

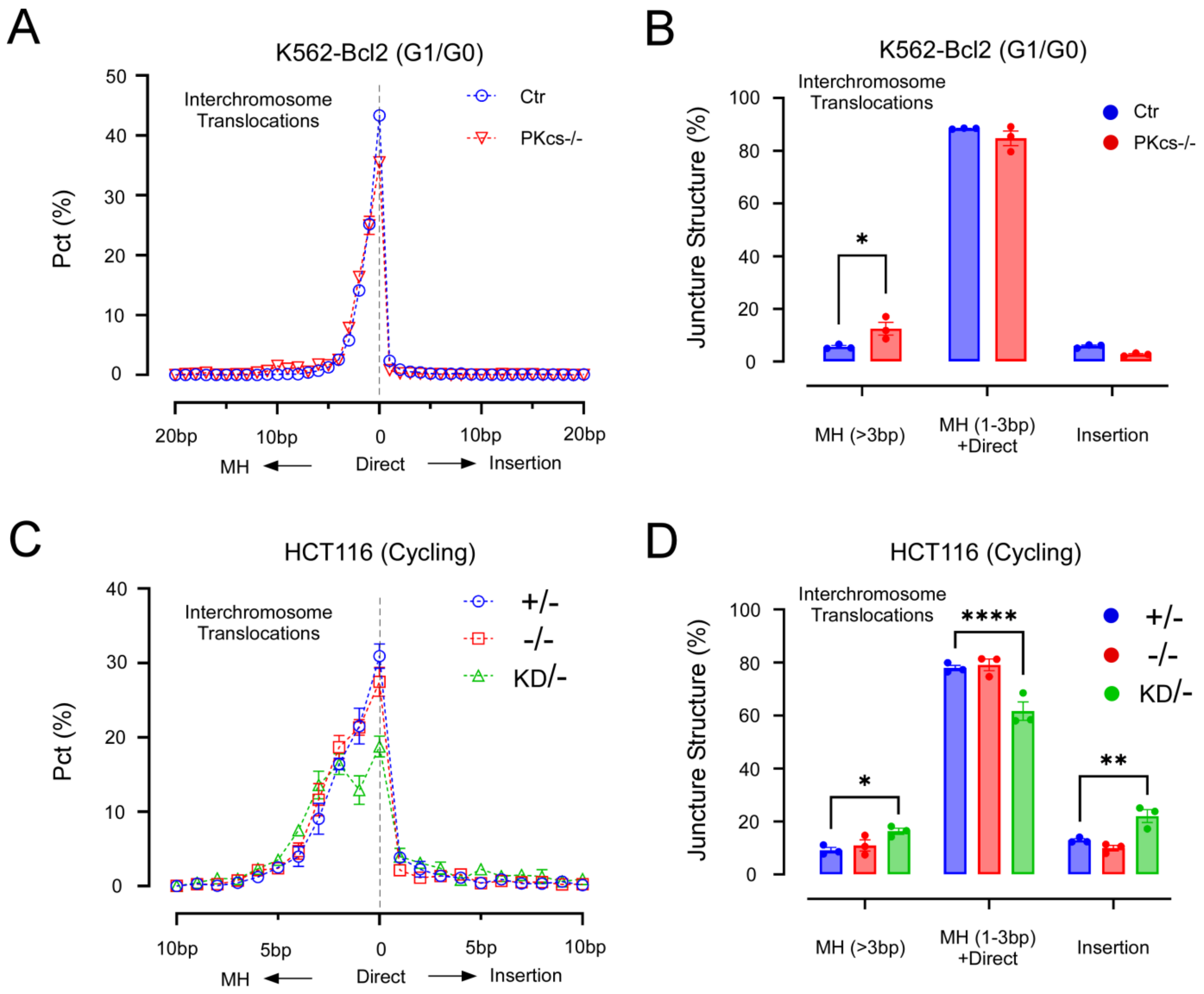


**Supplementary Figure S9. Impact of DNA-PKcs kinase domain alterations on MMEJ-mediated interchromosome translocations.** (**A**) Quantification of repair patterns in interchromosome translocations within a 20 bp window, including microhomology (MH), insertion, and direct repair. The data is presented as percentages (Pct, %). (**B**) The junction structures in (A) were divided into three subgroups, MH (>3bp), MH (1-3bp) plus Direct and Insertion, plotted and analyzed by two-way ANOVA plus Dunnett's multiple comparisons test with *p< 0.05. (**C**) Same as (A) but for HCT116 cycling cells with gene editing including *DNA-PKcs*$^{+/-}$, *DNA-PKcs*$^{-/-}$ and *DNA-PKcs*$^{KD/-}$ and ±10 bp window. (**D**) Same as (B) but for HCT116 cycling cells. All experiments were independently repeated three times (n=3), and the standard error of the mean (SEM) is indicated.

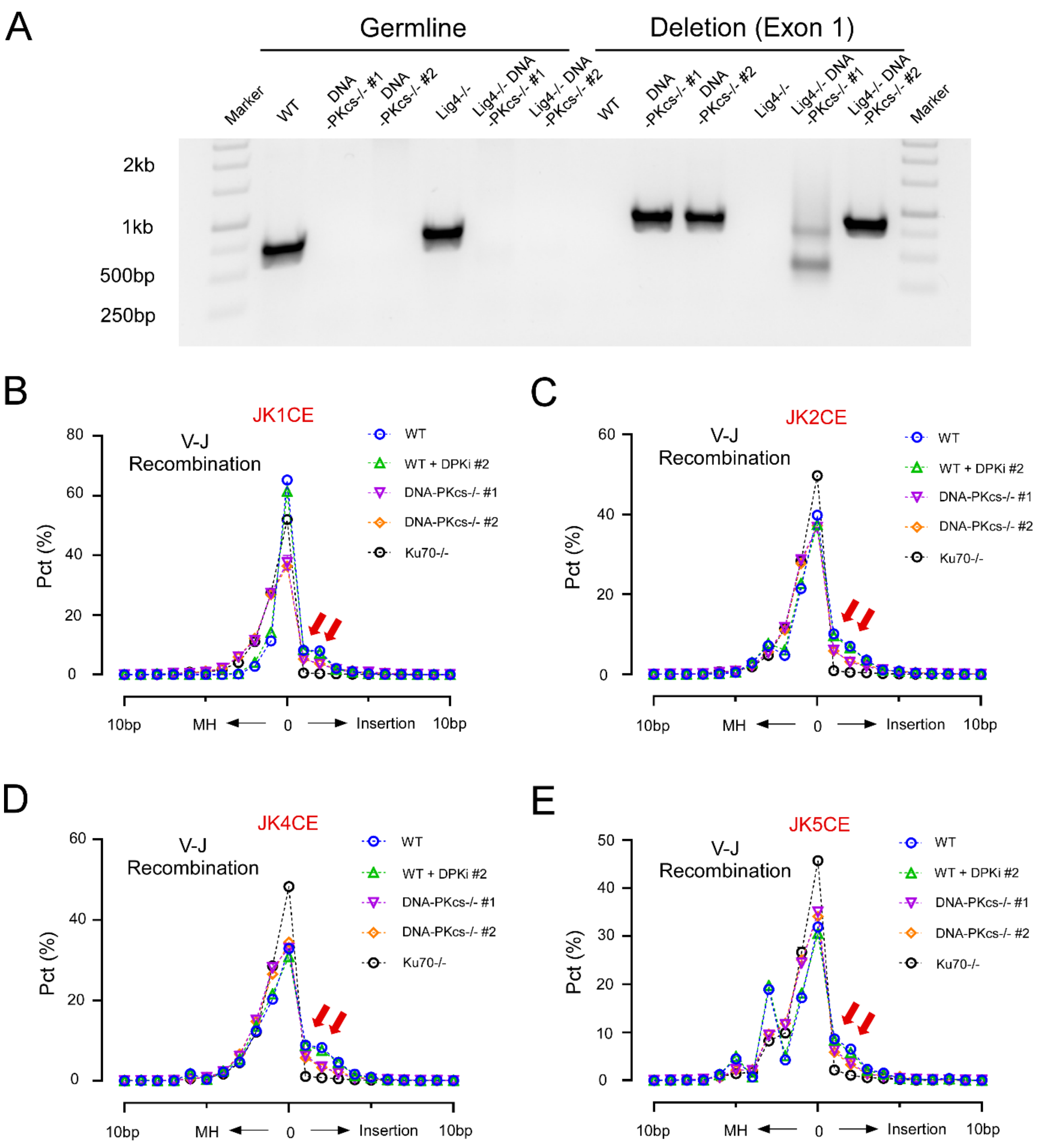


**Supplementary Figure S10. Reduced insertion rates in V-J recombination upon DNA-PKcs inhibition, deletion, and *Ku70*$^{-/-}$ in IgK antigen locus of vAbl ProB cells.** (**A**) PCR genotyping to confirm DNA-PKcs deletion (exon 1) in the *v-Abl* cell lines including *DNA-PKcs*$^{-/-}$ #1/2 and *Lig4*$^{-/-}$ *DNA-PKcs*$^{-/-}$ #1/2 while the *WT* and *Lig4*$^{-/-}$ cell lines served as control. Paired primers mDPK-seq-MF/MR and mDPK-seq-F/R (see Table S7) were used to detect the germline bands and deletional bands, respectively. (**B-E**) Repair pattern distribution of junctions obtained by HTGTS with JK1CE (**B**), JK2CE (**C**), JK4CE (**D**) and JK4CE (**E**) baits, including microhomology (MH), insertion, and direct repair, shown in a histogram with a 20 bp window in percentage (Pct, %). Red arrows indicate the drops in 1-2 bp insertion in conditions of DNA-PKcs inhibition (*WT* + DPKi #1/2), deletion (*DNA-PKcs*$^{-/-}$ #1/2), and *Ku70*$^{-/-}$, respectively. All experiments were independently repeated three times, and the standard error of the mean (SEM) is provided.

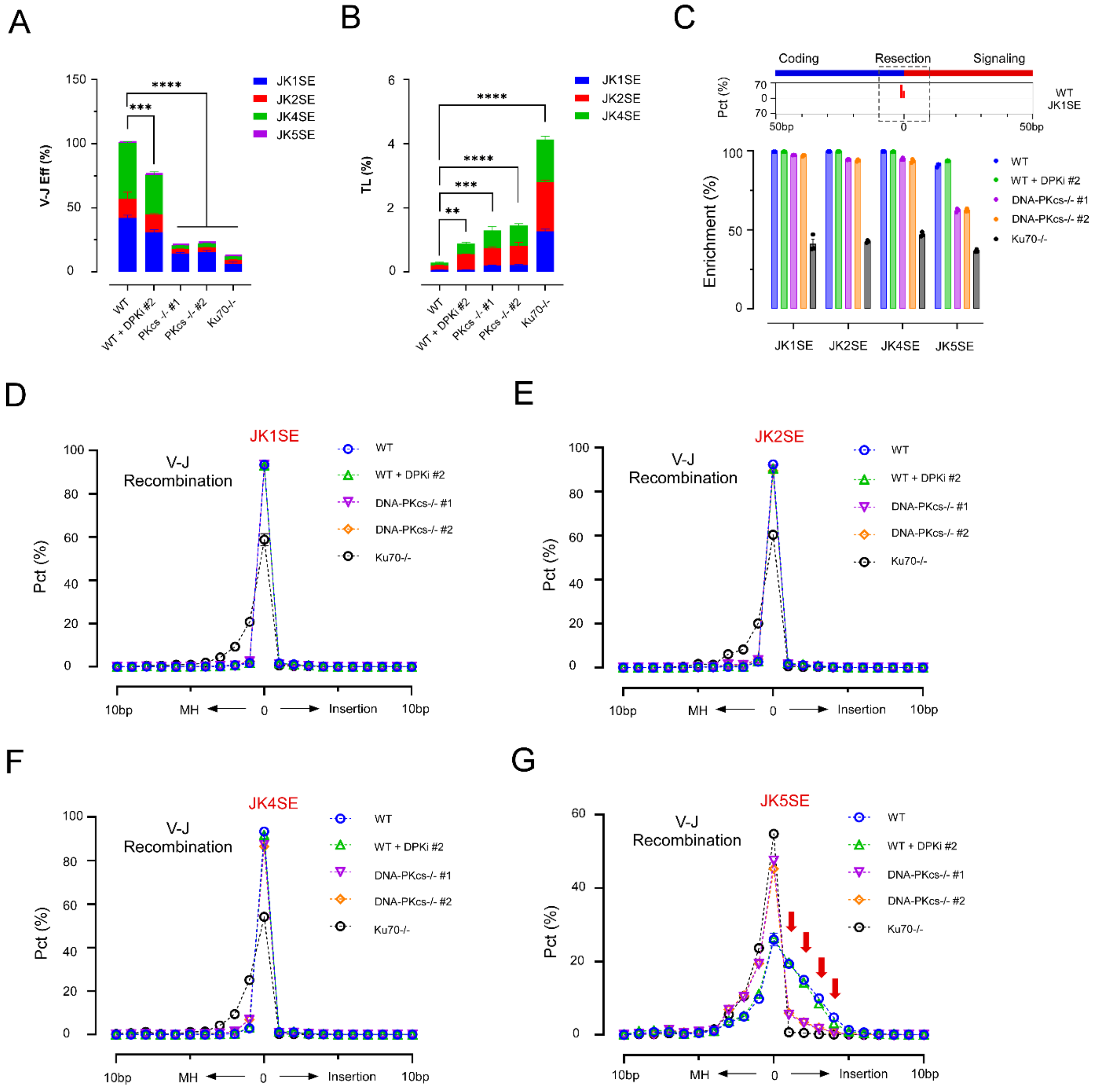


**Supplementary Figure S11. Impact of DNA-PKcs on V-J coding end recombination in murine IgK antigen locus with Jk signaling end (SE) bait.** (**A**) V-J recombination efficiency (V-J Eff) is shown as the number of V-J junctions normalized by the number of total reads in each library, using four functional Jk signaling end baits (JK1SE in blue, JK2SE in red, JK4SE in green, and JK5SE in magenta), and the results are presented as stacked bars. The combined values of *WT* (101.23 ± 4.28%, n=3), *WT* + DPKi #1 (104.34 ± 1.04%, n=3), *WT* + DPKi #2 (76.28 ± 4.54%, n=3), *DNA-PKcs*$^{-/-}$ #1 (20.71 ± 0.94%, n=3), *DNA-PKcs*$^{-/-}$ #2 (22.34 ± 1.32%, n=3), and *Ku70*$^{-/-}$ (12.63 ± 1.43%, n=3) were evaluated by ordinary one-way ANOVA plus Dunnett's multiple comparisons test. (**B**) Inter-chromosomal translocation is normalized by the total junctions as TL rate. The results for all robust baits JK1SE, JK2SE and JK4SE are shown in stacked bars. The combined values of *WT* (0.29 ± 0.02%, n=3), *WT* + DPKi #1 (0.34 ± 0.01%, n=3), *WT* + DPKi #2 (0.89 ± 0.05%, n=3), *DNA-PKcs*$^{-/-}$ #1 (1.29 ± 0.15%, n=3), *DNA-PKcs*$^{-/-}$ #2 (1.46 ± 0.15%, n=3), and *Ku70*$^{-/-}$ (4.13 ± 0.07%, n=3) were evaluated by ordinary one-way ANOVA plus Dunnett's multiple comparisons test. (**C**) The junctions distributed along all V gene fragment broken sites of a WT library with JK1SE bait are aggregated and plotted in a 100 bp window. The broken site is at position 0, flanked by coding (blue) and signaling (red) sequences. The degree of resection is the distance between the junction and the broken site. The junctions are predominantly found in the signaling site (red bars) and are enriched in a 20 bp resection window in WT. The junction enrichment in this 20 bp resection window is quantified and shown to compare the degree of resections in all conditions. (**D-G**) The repair pattern distribution of data with JK1SE (**D**), JK2SE (**E**), JK4SE (**F**), and JK5SE (**G**) baits, including microhomology (MH), insertion, and direct repair, in a 20 bp window in percentage (Pct, %). No detectable changes for all conditions except the *Ku70*$^{-/-}$ in panels (**D-F**). Significant shifts from insertion to direct repair were observed in conditions *DNA-PKcs*$^{-/-}$ #1/2 and *Ku70*$^{-/-}$ (red arrows) comparing with the WT in panel (**G**). All experiments were independently repeated three times, and the standard error of the mean (SEM) is provided with $^{**}p<0.01$, $^{***}p<0.001$ and $^{****}p<0.0001$.

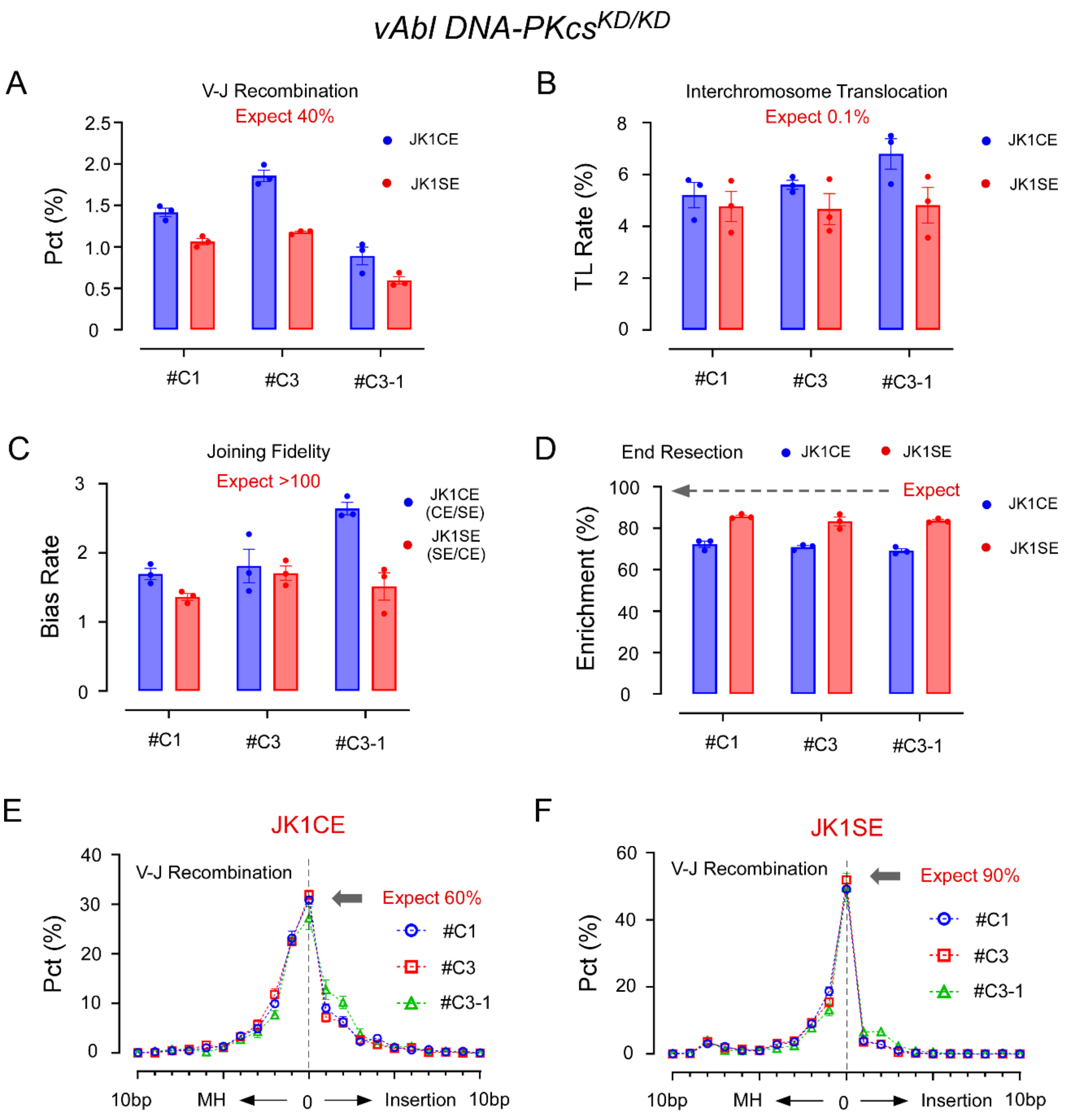


**Supplementary Figure S12. The impact of catalytically dead DNA-PKcs (*DNA-PKcs*$^{KD/KD}$) on V(D)J recombination in *vAbl* cells.** (**A**) The V-J recombination efficiency of three *vAbl DNA-PKcs KD* cells #C1, #C3 and #C3-1 obtained by JK1CE (blue bar) and JK1SE (red bar) baits, respectively. Note, the *WT vAbl* cells generated 40% V-J recombination (expect). (**B**) Same as (A) but for the interchromosome translocation, TL rate. (**C**) Same as (A) but for the joining fidelity with bias rate of more than 100 folds in *WT*. (**D**) Same as (A) but for the end resection, in contrast with expected nearly 100% junctions (arrow with dash line) fall within the ±10 bp window in *WT*. (**E**) The junction structure of V-J recombination obtained by JK1CE bait in three DNA-PKcs$^{KD/KD}$ cell lines. Note, direct repair (grey arrow) in *WT* was 60%. (**F**) Same as (E) but for the JK1SE bait. All experiments were independently repeated three times, and the standard error of the mean (SEM) is provided.

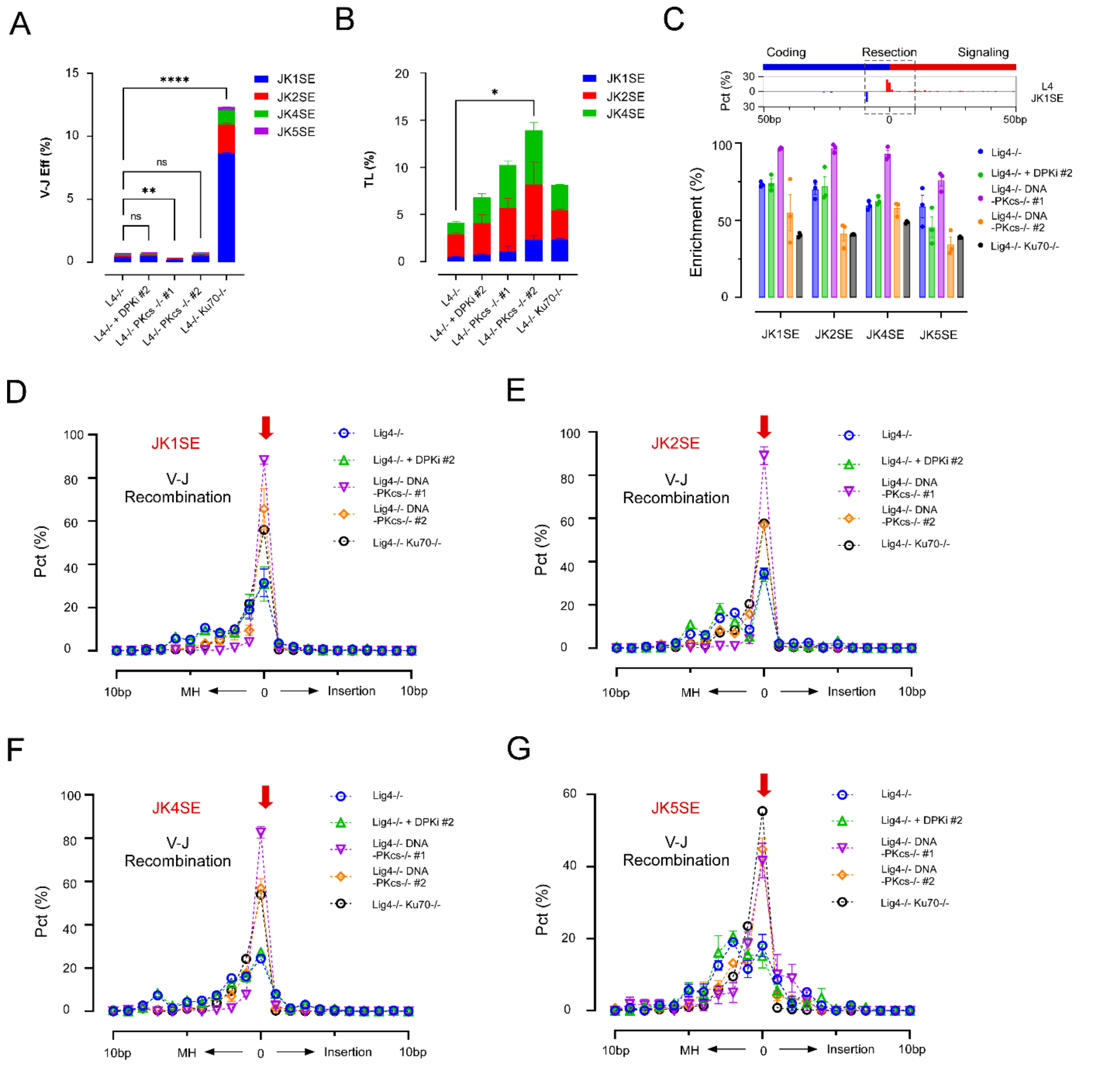


**Supplementary Figure S13. Impact of DNA-PKcs on V-J coding ends recombination in Ligase 4 deficient (*Lig4*$^{-/-}$) murine IgK antigen locus with Jκ signaling end (SE) bait.** (**A**) V-J recombination efficiency (V-J Eff) were systematically studied using four functional Jκ coding end baits (JK1SE in blue, JK2SE in red, JK4SE in green, and JK5SE in magenta), and the results are shown in stacked bars. The combined values of *Lig4*$^{-/-}$ (0.59 ± 0.01%, n=3), *Lig4*$^{-/-}$ + DPKi #1 (0.69 ± 0.03%, n=3), *Lig4*$^{-/-}$ + DPKi #2 (0.65 ± 0.01%, n=3), *Lig4*$^{-/-}$ + *DNA-PKcs*$^{-/-}$ #1 (0.20 ± 0.05%, n=3), *Lig4*$^{-/-}$ + *DNA-PKcs*$^{-/-}$ #2 (0.62 ± 0.12%, n=3), and *Lig4*$^{-/-}$ + *Ku70*$^{-/-}$ (12.33 ± 0.12%, n=3) were evaluated by ordinary one-way ANOVA plus Dunnett's multiple comparisons test. (**B**) Normalized inter-chromosomal translocation, TL rate for all signaling baits except JK5SE are shown in stacked bars. The combined values of *Lig4*$^{-/-}$ (4.10 ± 0.18%, n=3), *Lig4*$^{-/-}$ + DPKi #1 (3.89 ± 0.36%, n=3), *Lig4*$^{-/-}$ + DPKi #2 (6.86 ± 1.15%, n=3), *Lig4*$^{-/-}$ + *DNA-PKcs*$^{-/-}$ #1 (10.25 ± 1.44%, n=3), *Lig4*$^{-/-}$ + *DNA-PKcs*$^{-/-}$ #2 (13.94 ± 2.92%, n=3), and *Lig4*$^{-/-}$ + *Ku70*$^{-/-}$ (8.14 ± 0.18%, n=3) were evaluated by ordinary one-way ANOVA plus Dunnett's multiple comparisons test. (**C**) The junctions distributed along all V gene fragment broken sites of a *Lig4*$^{-/-}$ library with JK1SE bait are aggregated and plotted in a 100 bp window. The broken site is at position 0, flanked by coding (blue) and signaling (red) sequences. The degree of resection is the distance between the junction and the broken site. The junctions appeared in the coding site (blue bars) as well as signaling site (red bars) in L4$^{-/-}$. To compare the degree of resections in all conditions, the junction enrichment is quantified and shown. (**D-G**) The repair pattern distribution of data with JK1SE bait, including MH, insertion, and direct repair, in a 20 bp window in percentage (Pct, %). Red arrows indicate the shift from MH to direct repair in conditions of further *DNA-PKcs*$^{-/-}$ (#1/2) and *Ku70*$^{-/-}$. All experiments were independently repeated three times, and SEM are provided: *p<0.05, **p<0.01, ****p<0.0001.

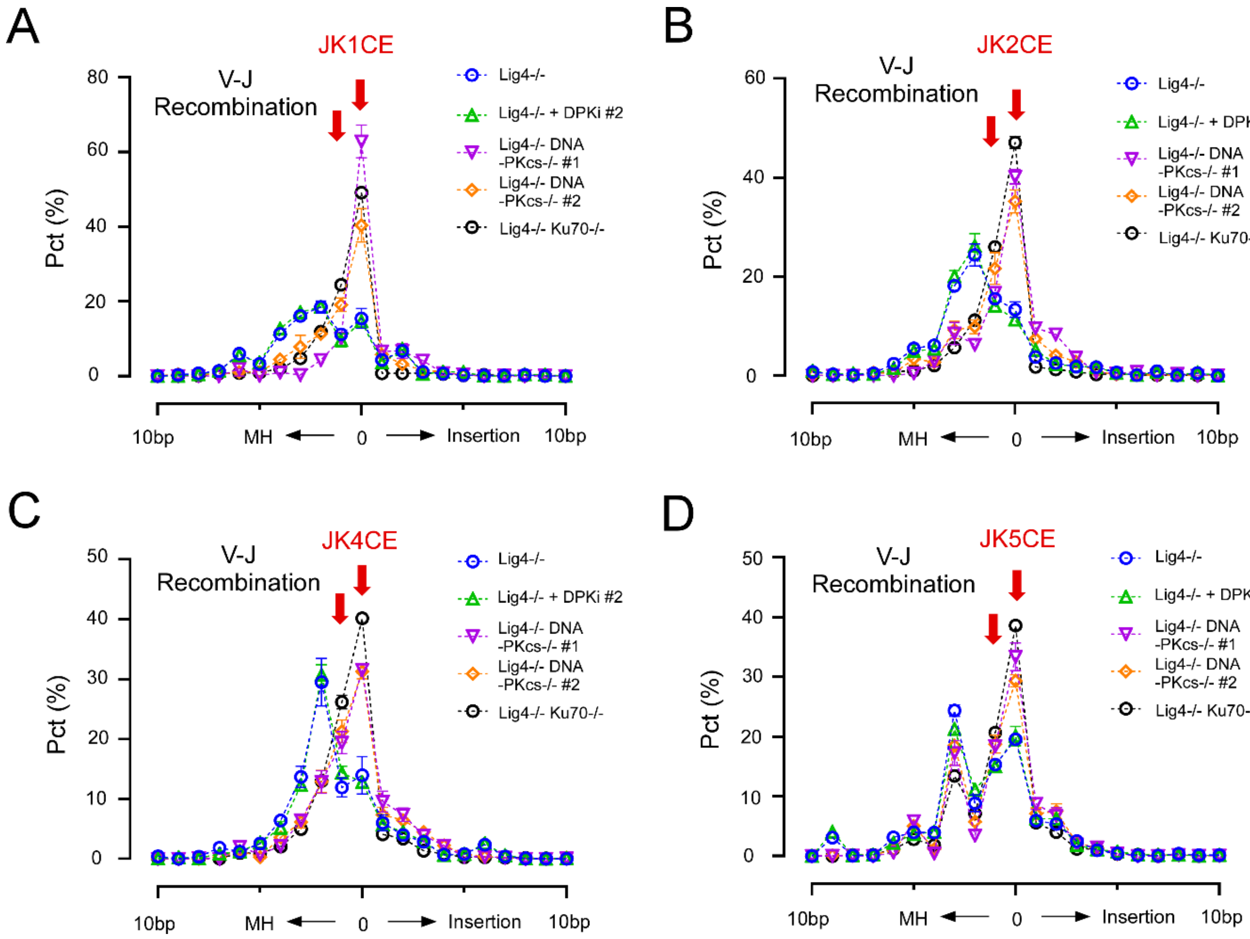


**Supplementary Figure S14. Enhanced direct repair rates in V-J recombination upon DNA-PKcs inhibition, deletion, and Ku70$^{-/-}$ in the context of Ligase 4 deletion (*Lig4*$^{-/-}$) in vAbl ProB IgK antigen locus.** (**A-D**) Histograms showing the repair pattern distribution of junctions obtained by HTGTS with JK1CE (**A**), JK2CE (**B**), JK4CE (**C**) and JK5CE (**D**) baits, including microhomology (MH), insertion, and direct repair, presented in a 20 bp window as a percentage (Pct, %). Red arrows highlight the shift in peak from microhomology to direct repair in conditions of DNA-PKcs deletion (*Lig4*$^{-/-}$ *DNA-PKcs*$^{-/-}$ #1/2), and Ku70 deletion (*Lig4*$^{-/-}$ *Ku70*$^{-/-}$), respectively, compared to the control (*Lig4*$^{-/-}$). All experiments were independently repeated three times, and the standard error of the mean (SEM) is provided.
